# Polypharmacology-based kinome screen identifies new regulators of KSHV reactivation

**DOI:** 10.1101/2023.02.01.526589

**Authors:** Annabel T. Olson, Yuqi Kang, Anushka M. Ladha, Chuan Bian Lim, Michael Lagunoff, Taran S. Gujral, Adam P. Geballe

## Abstract

Kaposi’s sarcoma-associated herpesvirus (KSHV) causes several human diseases including Kaposi’s sarcoma (KS), a leading cause of cancer in Africa and in patients with AIDS. KS tumor cells harbor KSHV predominantly in a latent form, while typically <5% contain lytic replicating virus. Because both latent and lytic stages likely contribute to cancer initiation and progression, continued dissection of host regulators of this biological switch will provide insights into fundamental pathways controlling the KSHV life cycle and related disease pathogenesis. Several cellular protein kinases have been reported to promote or restrict KSHV reactivation, but our knowledge of these signaling mediators and pathways is incomplete. We employed a polypharmacology-based kinome screen to identifiy specific kinases that regulate KSHV reactivation. Those identified by the screen and validated by knockdown experiments included several kinases that enhance lytic reactivation: ERBB2 (HER2 or *neu*), ERBB3 (HER3), ERBB4 (HER4), MKNK2 (MNK2), ITK, TEC, and DSTYK (RIPK5). Conversely, ERBB1 (EGFR1 or HER1), MKNK1 (MNK1) and FRK (PTK5) were found to promote the maintenance of latency. Mechanistic characterization of ERBB2 pro-lytic functions revealed a signaling connection between ERBB2 and the activation of CREB1, a transcription factor that drives KSHV lytic gene expression. These studies provided a proof-of-principle application of a polypharmacology-based kinome screen for the study of KSHV reactivation and enabled the discovery of both kinase inhibitors and specific kinases that regulate the KSHV latent-to-lytic replication switch.

**Author Summary:** Kaposi’s sarcoma-associated herpesvirus (KSHV) causes Kaposi’s sarcoma, a cancer particularly prevalent in Africa. In cancer cells, the virus persists in a quiescent form called latency, in which only a few viral genes are made. Periodically, the virus switches into an active replicative cycle in which most of the viral genes are made and new virus is produced. What controls the switch from latency to active replication is not well understood, but cellular kinases, enzymes that control many cellular processes, have been implicated. Using a cell culture model of KSHV reactivation along with an innovative screening method that probes the effects of many cellular kinases simultaneously, we identified drugs that significantly limit KSHV reactivation, as well as specific kinases that either enhance or restrict KSHV replicative cycle. Among these were the ERBB kinases which are known to regulate growth of cancer cells. Understanding how these and other kinases contribute to the switch leading to production of more infectious virus helps us understand the mediators and mechanisms of KSHV diseases. Additionally, because kinase inhibitors are proving to be effective for treating other diseases including some cancers, identifying ones that restrict KSHV replicative cycle may lead to new approaches to treating KSHV-related diseases.

## Introduction

Kaposi’s sarcoma-associated herpesvirus (KSHV) is the etiologic agent of Kaposi’s sarcoma (KS), a leading cause of cancer in Africa and a substantial health concern for AIDS patients worldwide [1–3]. KSHV causes three other less prevalent diseases: primary effusion lymphoma in B cells, multicentric Castleman disease, and a KSHV inflammatory cytokine syndrome. The main proliferating tumor cell of KS is the spindle cell. In KS spindle cells, the virus exists predominantly in a latent state in which only a few of its ∼90 genes are expressed. Approximately 5% of spindle cells express markers of the lytic replicative cycle [4–7] representing a relatively infrequent switch from latency to lytic replication in tumors. While considerable effort has been devoted to characterizing cellular and viral factors that support the maintenance of latency or the induction of lytic replication, our knowledge of the complex signaling involved in this replicative switch remains incomplete. Because many latent and lytic KSHV genes have oncogenic properties and are involved in disease progression [4, 8–13], understanding the regulators of this replicative switch is of fundamental importance for understanding KSHV disease pathogenesis and has potential relevance for new therapeutic interventions..

The human kinome comprises 518 protein kinases known to regulate myriad host and viral processes, including KSHV latency and the switch to lytic replication [14–16]. Due to the essential regulatory roles of kinases, dysregulation of their catalytic activity causes many types of cancers and other diseases. Viruses also usurp cellular kinases or encode their own kinases to modulate the signaling of the host cell to promote specific virus lifecycle stages or replicative functions. For KSHV, both the virus-encoded kinase, ORF36 [17], and cellular kinases are necessary for lytic replication [18–22]. Prior reports identified several kinases with roles in KSHV reactivation using various screening approaches, including a kinase cDNA overexpression screen [23], phospho-site antibody microarray [18], and proteome analysis [24] following KSHV primary infection or after induction of lytic replication. From these and other studies, fewer than a dozen kinases have been validated as aiding latency or facilitating reactivation [18, 24, 25]. By completing a more comprehensive investigation of protein kinase regulators of KSHV reactivation, we might identify FDA-approved kinase inhibitors that could be repurposed with the aim of reducing KSHV reservoirs and/or treating KSHV-associated cancers and lymphoproliferative diseases.

Recently, kinase centric polypharmacology-based screens have been developed to identify both kinase inhibitors and their targeted kinases that regulate cell death, cancer cell migration and other cancer cell phenotypes [26–28]. These screens have also been used in *Plasmodium* infected cells and to evaluate kinase roles during virus-induced cytokine production [29, 30]. This innovative approach employs broadly acting kinase inhibitors as tools that exploit built-in redundancy from their shared kinase targets, when used in Kinase Inhibitor Regularization (KiR) analyses [26, 27, 31]. Specifically, the polypharmacology-based kinome screen and KiR platform uses a small set of computationally-derived kinase inhibitors to restrict the catalytic activity of multiple endogenously expressed kinases. Data derived from testing the phenotypic effects of these inhibitors, coupled with known drug specificities and potencies for each kinase target, allows for a network-based, machine-learning analysis that initially predicts the impact of untested kinase inhibitors. Refinements by iterative screening of additional kinase inhibitors curates with high accuracy, single kinases predicted to have significant regulatory potential for the system evaluated. This method is attractive compared to alternative kinome screening methods due to the high-throughput nature, built-in redundancy for enhanced accuracy, and two screen outputs, kinase inhibitors and specific kinases.

Herein, we describe the adaption of this kinome screening approach to study KSHV reactivation in an epithelial cell system commonly used to study KSHV reactivation. From this approach, we discovered two drugs, lestaurtinib and K252a, as potent kinase inhibitors of induced lytic replication and an initial six kinases not previously associated with reactivation. Among these predicted kinases, MKNK2 and ERBB4 had the greatest pro-lytic phenotypes. MKNK2 (MNK2) and the closely related MKNK1 (MNK1) are both mitogen-activated kinases and are the only kinases known to phosphorylate eIF4E [32].The epidermal growth factor receptor kinase ERBB4 has three other family members, all of which regulate numerous cell-signaling pathways including transcription, translation, cell survival, and cell growth (Kumagai 2021; Burgess 2022). Evaluation of the other members of these kinase families revealed that ERBB2 (HER2 or *neu*) and ERBB3 are pro-lytic factors, like MKNK2 and ERBB4, while MKNK1 and ERBB1 (EGFR) have opposite, pro-latent effects. Downstream signaling characterization of ERBB2 during KSHV reactivation revealed a new connection between ERBB2 and activation of CREB1, a transcription factor known to activat KSHV lytic gene expression [18]. Based on our findings, we propose a model in which ERBB1:ERBB2 heterodimer signals to promote latency. Next, induction of lytic replication turns on ERBB3 expression, providing a higher affinity-binding partner for ERBB2 and steals ERBB2 away from ERBB1. Due to similar lytic-promoting phentoypes of ERBB2 and ERBB3, we predict that this newly formed heterodimer signals through CREB1 to activate lytic gene transcription. These findings provide initial insights into how KSHV latent-to-lytic replication switch can be regulated, by interfering with ERBB signaling, and suggest potential utility of manipulating signaling from these plasma membrane receptors as a new therapeutic approach.

## Results

### Generation of a recombinant KSHV with infection and lytic replication indicators

To enable precise measurement of the initial transition from KSHV latent-to-lytic replication, we constructed a new recombinant virus called lytic replication indicator KSHV (KSHV^LRI^). This virus is derived from the KSHV bacterial artificial chromosome 16 (BAC16) that expresses GFP constitutively, enabling identification of infected cells. For detecting lytic replication, we introduced an expression cassette containing the KSHV polyadenylated long non-coding RNA promoter (PrPAN) driving expression of a streptavidin-binding peptide fused to a truncated low-affinity nerve growth factor receptor (SBP-ΔLNGFR) and nuclear localized mCherry (mCherry-NLS) (Fig 1A). The coding regions for these two genes are separated by a 2A self-cleaving peptide (P2A) sequence. The SBP-ΔLNGFR was designed for selection by streptavidin binding of cells undergoing lytic replication. The NLS on mCherry enables accurate, high-throughput quantification of individual cells containing lytic replicating virus by fluorescence imaging. We introduced KSHV^LRI^ into iSLK cells, a human renal carcinoma epithelial cell line that encodes doxycycline (DOX)-inducible KSHV replication and transcription activator (RTA). As expected, the KSHV^LRI^ infected cells expressed GFP (Fig 1B) and LANA (Figs S1A and S1B), and addition of lytic inducing agents, DOX or DOX plus sodium butyrate (NaB), resulted in mCherry-NLS (Fig 1B) and SBP-ΔLNGFR (Figs S1C and S1D) expression. Importantly, the engineered fluorescent constructs allowed us to monitor total KHSV^LRI^ infected cells by GFP area per image and virus reactivation by mCherry-NLS positive cells per image in infected iSLK cells using high-throughput, quantitative fluorescence imaging.

**Fig 1:**
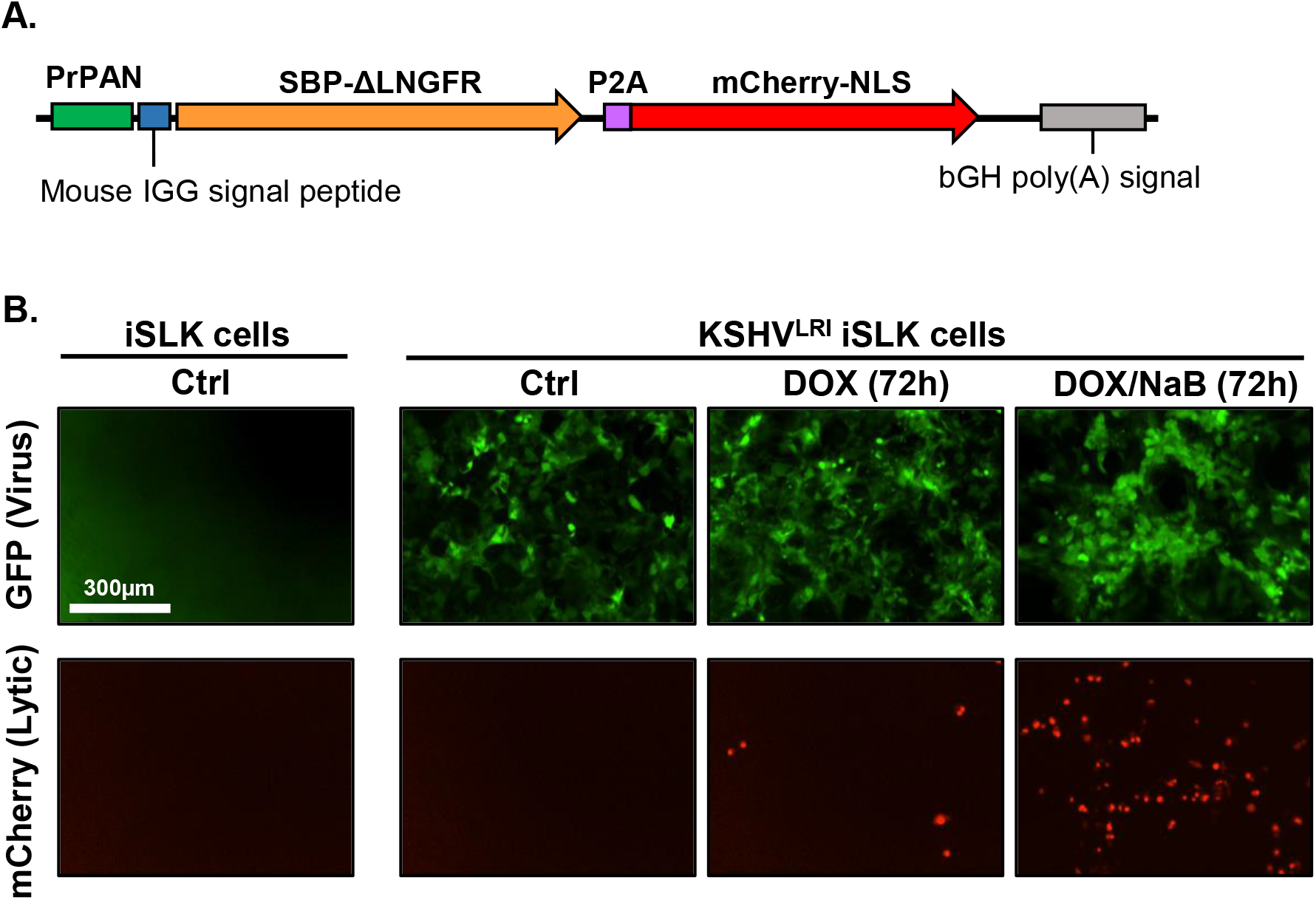
Characterization of KSHV dual lytic replication indicator virus (KSHV^LRI^) reactivation in iSLK cells. **(A)** Diagram of DNA cassette inserted into KSHV BAC16 genome to generate the KSHV^LRI^ recombinant. The KSHV *PAN* promoter (PrPAN) drives expression of the streptavidin-binding peptide fused to a truncated low-affinity nerve growth factor receptor (SBP-ΔLNGFR), a P2A self-cleaving peptide sequence, and then mCherry containing a nuclear localizing signal (mCherry-NLS). **(B)** Fluorescence images of uninfected iSLK and KSHV^LRI^ infected cells collected using an Incucyte Imaging System. Infected cells express the virus encoded *GFP* under control of the cellular EF-1α promoter (top row). mCherry-nls (bottom row) detected after inducing lytic replication with 1µg/ml DOX (middle column) or DOX plus 1mM NaB (right column) for 72h.

### Polypharmacology-based kinome screen for enhancers or repressors of KSHV reactivation

To identify kinases important for KSHV reactivation, we employed a polypharmacology-based kinome screen using the kinase inhibitor regularization (KiR) pipeline as previously described [31]. First, we incubated KSHV^LRI^ latently infected iSLK cells with the vehicle (DMSO) control or each of the 29 computationally-selected kinase inhibitors (KI 1-29) at four concentrations (31nM, 125nM, 500nM, 2µM) and concurrently induced lytic replication by adding DOX plus NaB to the cell media (Fig 2A). Collectively, these 29 KIs targeted a broad range of kinases (296 out of 360 kinase profiled). At 72h post-treatment, we quantified KSHV reactivation by counting mCherry-NLS positive cells per image and cell survival by measuring GFP area per image using the Incucyte Imaging System. A representative dose-response curve for two kinase inhibitors (cmpd #1: CAS no. 252003-65-9; cmpd #2: CAS no. 1221153-14-5) that either enhanced or restricted KSHV reactivation at 500nM concentration is shown in Fig 2B. We found that seven of the KIs were toxic to cells under these conditions and these were removed from further analyses. Data from the remaining 22 KIs at 500nM with DOX plus NaB provided the initial “training set” for machine learning-based analyses [33] with a range of 6 – 30% total cells reactivated. For the DMSO control, 20% total cells were reactivated and set to 100 (Fig 2C). While there were changes in the induction of lytic replication following treatment with only the KIs (without DOX or NaB), the range of lytic replication (0 – 0.06% total cells) was too small for the KiR analysis to generate kinase inhibitor predictions (Fig S2). These data suggest that broad kinase inhibition alone cannot efficiently activate the switch from latency to lytic replication in this system, but following RTA expression and release of some epigenetic constraints by NaB, kinases do measurably regulate KSHV reactivation.

**Fig 2:**
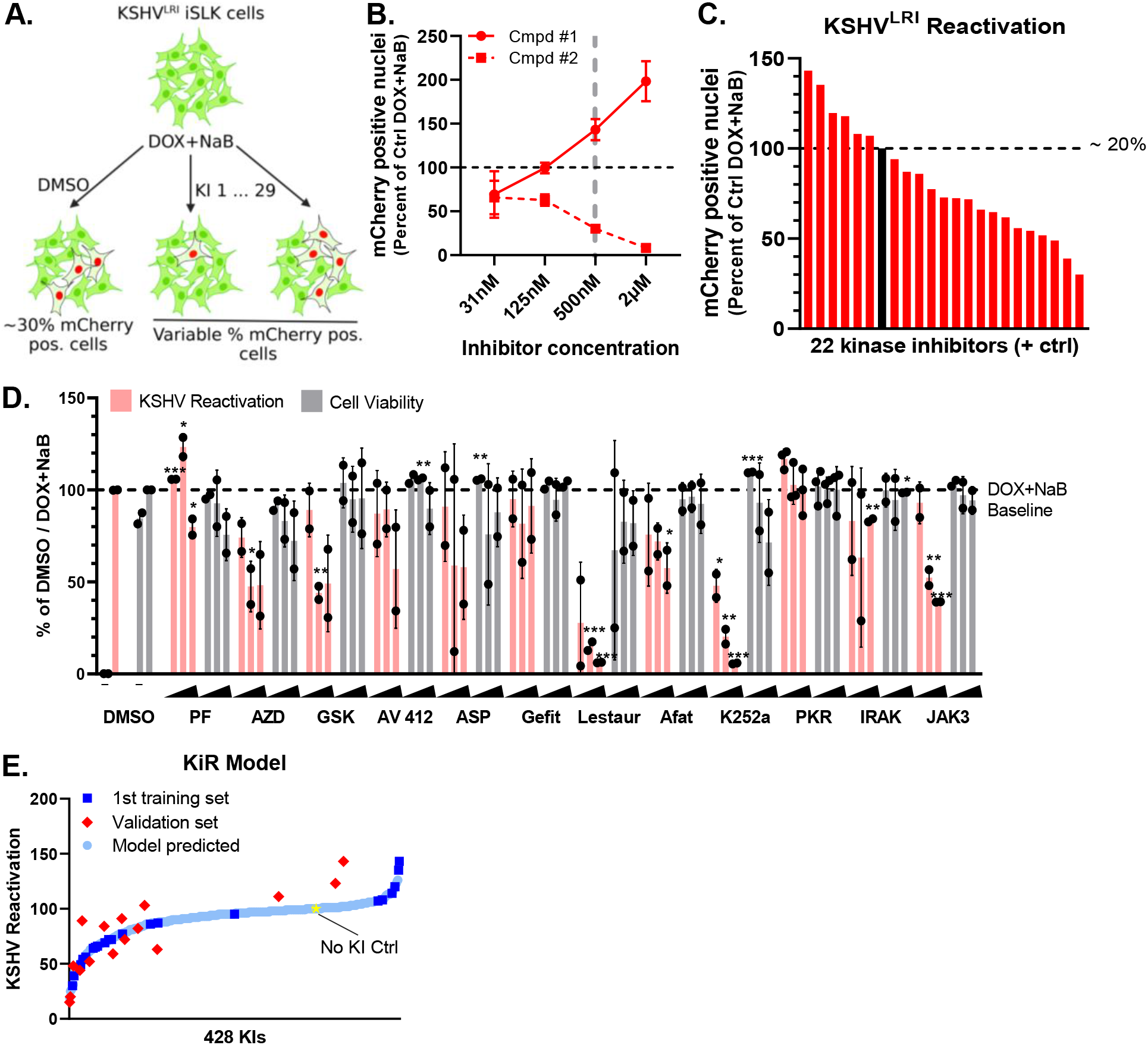
Polypharmacology-based kinome screen to identify kinases important for restriction or enhancement of KSHV lytic replication. **(A)** 29 selected kinase inhibitors (KIs) were tested at four concentrations (0.031, 0.125, 0.5, and 2uM) in KSHV^LRI^ latently infected iSLK cells in combination with KSHV lytic replication inducing agents. KSHV reactivation was measured by Incucyte Imaging System quantification of mCherry-NLS expression at 72h following treatment and provided the phenotypic data for the machine-learning analyses and prediction of kinases important for KSHV reactivation. **(B)** Data from testing two of these proprietary compounds illustrate dose-dependent increased or restricted KSHV reactivation. **(C)** KSHV^LRI^ reactivation phenotypes were obtained from 22 of the 29 selected KIs that did not cause cellular toxicity. Reactivation data for DOX plus NaB treatment alone (black bar and dotted black line set to 100, represents ∼20% total cells) or combined with KIs (red bars, 0.5 µM) were calculated as a percent of DOX plus NaB control. **(D)** Twelve additional KIs predicted from the initial model were tested for KSHV reactivation (pink bars) and cell viability (grey bars). The iSLK KSHV^LRI^ cells were untreated (“-” DMSO only) or treated with DOX plus NaB alone (DMSO) or in combination with KIs at 125nM, 500nM, or 2μM. Reactivation (pink bars) for each KI condition was measured by mCherry expression and calculated as a percent of DOX plus NaB with DMSO control. Cell viability (grey bars) was determined by cell confluence as a percent of DOX plus NaB with DMSO control. PF: PF-477736; AZD: AZD3463; GSK: GSK-650394; ASP: ASP-3026; Gefit: Gefitinib; Lestaur: Lestaurtinib; Afat: Afatinib; PKR: PKR Inhibitor; IRAK: IRAK1/4; JAK3: JAK3 Inhibitor VI. P-values for * ≤ 0.05, ** ≤ 0.01, *** ≤ 0.001. **(E)** The KiR model was developed by incorporating data on kinase inhibitors as the training set (C). This data was used to predict the response of potential kinase inhibitors. A subset of these predictions was then tested as a validation set (D), and the results were used to refine the model’s predictions. The No KI Ctrl (yellow star) represents the DOX plus NaB with DMSO control condition.

For the lytic inducing condition (DOX + NaB) that had a greater dynamic range for KSHV reactivation, we produced an initial KiR model from the 22 KI “training set” phenotype dataset and a separate dataset of > 400 KI’s effects on 298 recombinant human protein kinases [33]. Leave-one-out-cross validation (LOOCV) was used to evaluate the accuracy of the model. We then tested an additional 12 KIs predicted by the KiR model to impact KSHV reactivation (Fig 2D, Table S1). The resulting responses were iteratively included in the training set to improve the prediction accuracy of the model. Based on the final KiR model, the correlation between predicted and observed response was >0.8. The final list of 428 predicted kinase inhibitors is shown in Fig 2E and Table S2. Of note, two broadly acting kinase inhibitors, lestaurtinib and K252a, were predicted to be regulators of reactivation based on the initial KiR model. Following testing of these drugs (Fig 2D) and inclusion of the KSHV reactivation data into the final model, these two drugs remained at the top of the list of kinase inhibitors predicted to regulate KSHV reactivation (Table S2).

### Kinases validated as cellular regulators of KSHV reactivation in epithelial cells

Based on the KiR model, we selected 13 kinases with the highest average coefficient across all alpha values (Table S3) which corresponds to likely regulators of KSHV reactivation. We evaluated these predictions using siRNAs targeting each of these kinases. Individual depletion of six of these kinases significantly altered KSHV reactivation under lytic inducing conditions without having significant effects on cell viability (Fig 3). Knocking down ERBB4 (HER4), MKNK2 (MNK2), and DSTYK (RIPK5) as well as two TEC family kinases, ITK and TEC, reduced reactivation, indicating that these kinases are pro-lytic factors. In contrast, knocking down FRK (RAK or PTK5) caused a slight but statistically significant increase in reactivation, suggesting that it may be a pro-latency factor.

**Fig 3:**
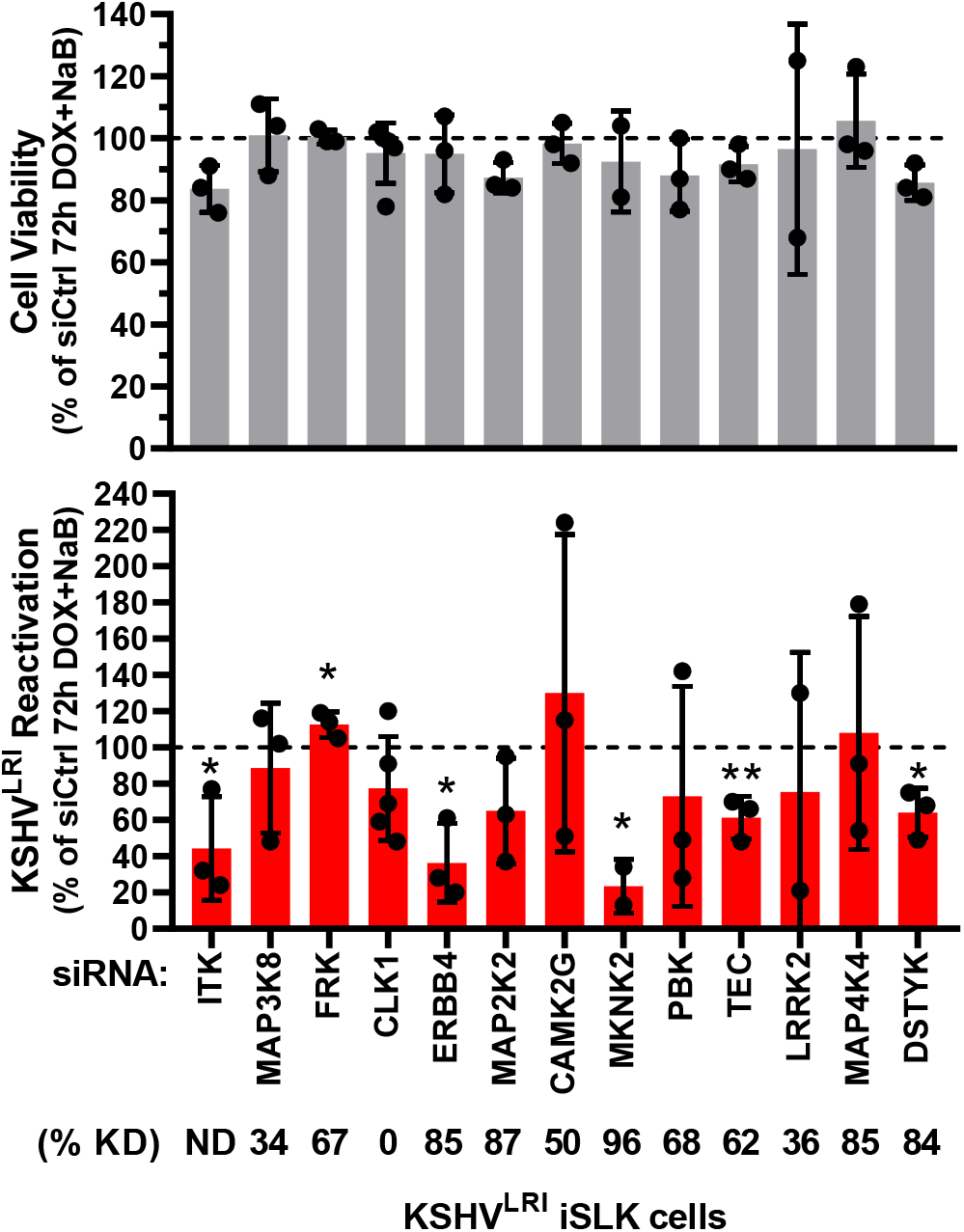
Validation of kinases predicted to regulate KSHV reactivation. Cell viability (grey bars) and KSHV reactivation (red bars) were measured for siRNA treated cells at 6-days post siRNA transfection and 3-days post DOX and NaB treatment. Control siRNA transfected cells treated with DOX plus NaB (dotted black line) was set to 100 and data for each kinase knockdown condition was calculated as a percent of the control. Kinase knockdown efficiencies at 3-days following siRNA transfection were determined before addition of lytic inducing drugs and graphed in Figs S3 and S4. For each knockdown, the efficiencies were averaged and listed below the corresponding kinase target as % KD. P-value * < 0.05 and ** < 0.01.

Our finding that knocking down ITK reduced reactivation (Fig 3) was surprising since we did not detect ITK expression by RT-qPCR in our latently infected cells. Consistent with our expression results, RNA-seq data from KSHV BAC16 latently infected iSLK cells showed very low ITK expression ([34], S4 Table). However, that study also revealed a >5-fold increase in ITK expression at 48h post lytic induction. The observation that knocking down another TEC family kinase (TEC) also restricted reactivation supports the conclusion that TEC kinase family signaling likely contributes to KSHV reactivation.

Because knocking down ERBB4 and MKNK2 had the strongest inhibitory effect on KSHV reactivation and both have other family members with shared substrates and overlapping signaling pathways, we tested the impact of knocking down these related kinases. As expected, ERBB2 and ERBB3 showed pro-lytic functions (Fig 4A) similarly to ERBB4. ERBB1 knock down had a similar trend but was not statistically significant. Intriguingly, MKNK1 had the opposite, pro-latent effect as compared to MKNK2 pro-lytic activity (Fig 4B). In these cells we showed that kinase knockdown was generally quite efficient by RT-qPCR assays (Figs S3 and S4) and that the siRNAs targeting ERBB or MKNK family members were specific for the individual target (Fig S4).

**Fig 4:**
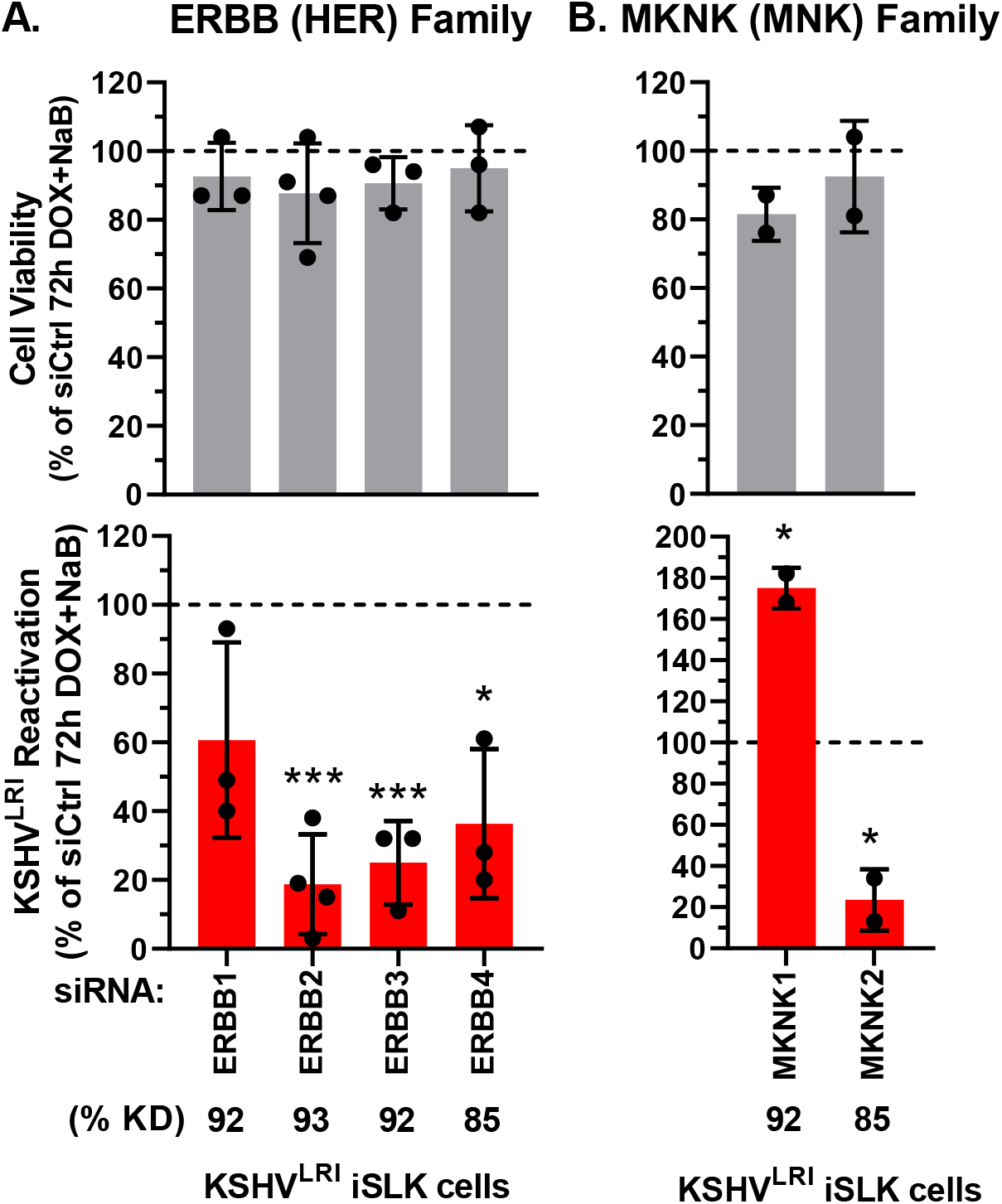
Effects of EBBB and MKNK family kinases on KSHV reactivation. Cell viability (grey bars) and KSHV reactivation (red bars) were measured at 6-days post transfection with siRNAs targeting **(A)** ERBB or **(B)** MKNK family kinases and 3-days post DOX and NaB treatment. Control siRNA transfected cells treated with DOX plus NaB (dotted black line) was set to 100 and data for each kinase knockdown condition was calculated as a percent of the control. Kinase knockdown efficiencies at 3-days following siRNA transfection were determined before addition of lytic inducing drugs and graphed in Fig S4. For each knockdown, the efficiencies were averaged and listed below the corresponding kinase target as % KD. P-value * < 0.05 and *** < 0.001.

Together, these data identify MKNK1, MKNK2, ERBB2, ERBB3, ERBB4, ITK, TEC, DSTYK, and FRK as kinases involved in KSHV reactivation and have pro-lytic functions except for MKNK1 and FRK which were pro-latency factors. One prior study reported that a kinase inhibitor, that moderately restricts MKNK1 and MKNK2 catalytic activity, reduced lytic replication [35] but otherwise, to the best of our knowledge, all the kinases we validated represent uncharacterized regulators of KSHV.

### ERBB2 and ERBB3 promote while ERBB1 restricts KSHV lytic gene expression

We prioritized further study of the effects of ERBB kinases on reactivation, as three of these kinases shared pro-lytic properties. First, we quantified the impact of depleting each ERBB on the transcription of the lytic *PrPAN-mCherry* (an early gene), *ORF10* (a late gene), and *K8.1* (a late gene) upon incubation with lytic inducing agents. To our surprise, knockdown of ERBB1 significantly increased lytic *PrPAN-mCherry* and *ORF10* transcript abundance and had a similarly trend for late lytic *K8.1* expression (Fig 5). These data do not correlate with the KSHV reactivation data (Fig 4) and suggest that ERBB1 may signal through at least two distict pathways to regulate KSHV reactivation. ERBB2 and ERBB3 knockdown significantly reduced both *PrPAN-mCherry* and *ORF10* expression in lytically induced cells (Figs 5A and 5B), consistent with the mCherry phenotype (Fig 4A). Expresson of *K8.1* was restricted by either ERBB2 or ERBB3 knockdown, but these changes were not statistically significant (Fig 5C). These data support roles for ERBB2 and ERBB3 as promoters of KSHV lytic gene expression while ERBB1 promotes latency by restricting lytic gene expression.

**Fig 5:**
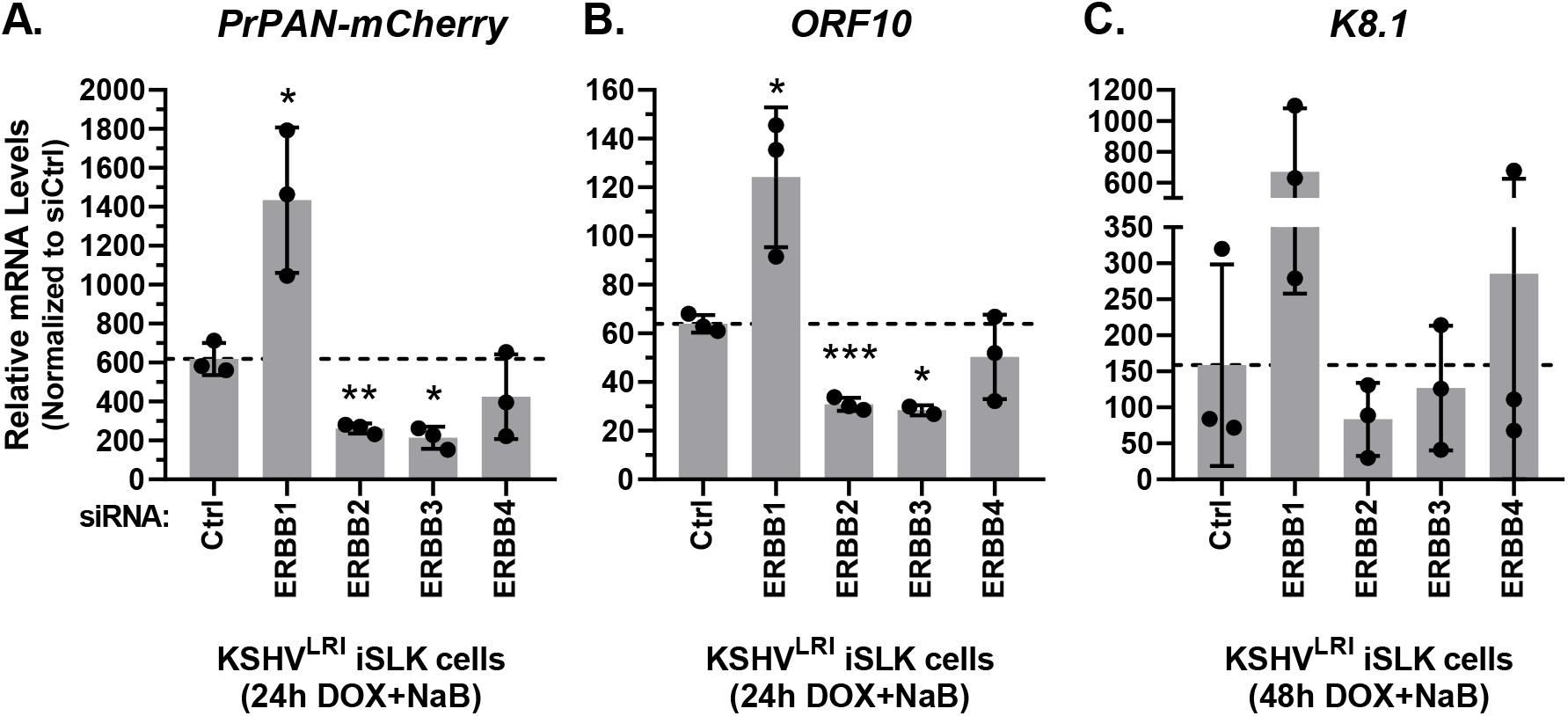
Effects of ERBB family member depletion on lytic gene transcript accumulation. KSHV^LRI^ latently infected iSLK cells were transfected with siRNA control or siRNAs targeting individual ERBB family kinases and then 3-days later treated with DOX plus NaB for 24h or 48h. Transcript levels for **(A)** *PrPAN-mCherry* **(B)** *ORF10* and **(C)** *K8.1* were quantified using target specific primers and RT-qPCR of total RNA.

### ERBB2 activation of ERBB1 signaling is disrupted during KSHV latent-to-lytic replication switch

Because ERBB2 is the preferential binding partner of all other ERBBs [36] and ERBB3 cannot signal alone [37], we focused our attention on ERBB2 signaling during KSHV reactivation. The ERBB signaling cascades are complex [38, 39]. Therefore, we measured activation of a panel of substrates (Fig 6A) using a high-throughput, reverse-phase protein array approach under both latent and induced lytic replication conditions in control and ERBB2 depleted cells. Depletion of ERBB2 in latently infected cells significantly attenuated the activation of ERBB1, as measured by phosphorylation at Tyr^1173^ (Fig 6B). This result suggests that in latent cells, ERBB1:ERBB2 heterodimers form and ERBB2 transphosphroylates ERBB1 to activate downstream signaling. Treatment with lytic inducing agents (in control siRNA cells) attenuated ERBB1 phosphorylation to a similar extent as in latently infected cells with ERBB2 depletion. ERBB2 knockdown in cells treated with lytic inducing agents did not further reduce ERBB1 activation. These results suggest that the ERBB1:ERBB2 heterodimer is disrupted during induction of lytic replication and are consistent with our finding that ERBB1 is a pro-latent factor that restricts lytic gene expression (Fig. 5).

**Fig 6:**
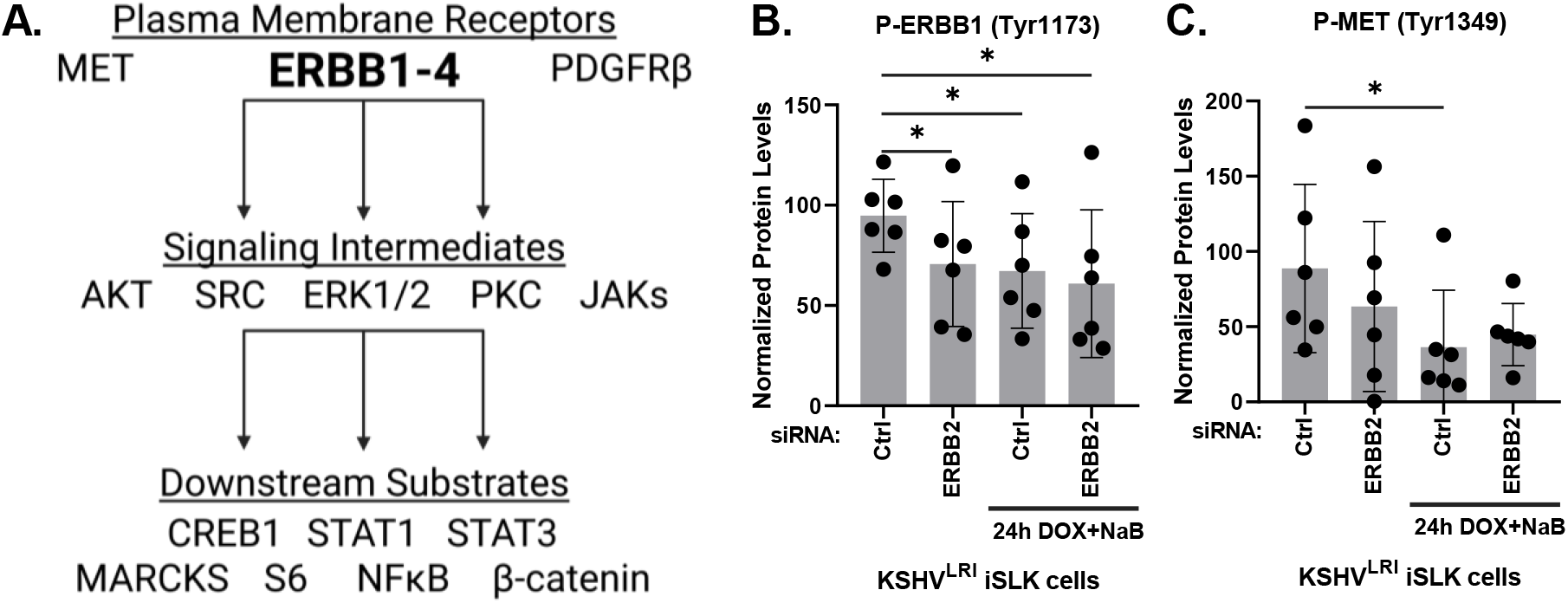
ERBB2 effects on ERBB1 and MET phosphorylation. **(A)** Diagram of selected proteins tested to elucidate ERBB2 signaling. **(B)** KSHV^LRI^ latently infected iSLK cells were transfected with siRNA control or siRNAs targeting ERBB2 and then 3-days later untreated or treated with DOX plus NaB for 24h. Cells were harvested, and protein lysates were analyzed using a reverse-phase protein array (RPPA) for ERBB1 phosphorylation at Tyr^1173^ or **(C)** for MET phosphorylation at Tyr^1349^. P-value * ≤ 0.05.

In our panel of substrates, we also evaluated phosphorylation of proteins involved in parallel signaling or crosstalk with EBBB2, as well as downstream signaling intermediates, and substrates of the selected signaling intermediates (Fig 6A). For proteins involved in crosstalk signaling, we measured the activation of plasma membrane receptors MET [40–42] and PDGFRβ [43] during lytic reactivation in control and ERBB2 knockdown cells. Similar to ERBB1, we found that MET activation decreased during lytic induction as compared to latent infection (Fig 6C), however, PDGFR-β was unchanged (Fig S5A). Furthermore, ERBB2 knockdown did not affect MET nor PDGFR-β activation, indicating that neither are regulated by ERBB2 during latency or reactivation.

In testing downstream signaling intermediates, we found that depletion of ERBB2 significantly reduced AKT activation during latency (Fig 7A). Activation of SRC and ERK1/2 showed a similar trend but was not statistically significant. Strikingly, the phosphorylation of all these kinases were similarly decreased following reactivation. Depleting ERBB2 during reactivation did little to alter these trends, although it did slightly increase ERK1/2 phosphorylation as compared to control lytically induced cells (Fig 7A). No significant trend was observed for pan PKC phosphorylation (Fig S5B). The reduced phosphorylation of AKT, SRC and ERK1/2 mimicked the decreased ERBB1 phosphorylation phenotypes and suggest that ERBB1:ERBB2 signaling through AKT and to a lessor degree SRC and ERK1/2 is ERBB2 dependent in latent cells and disrupted during early stages of lytic replcation.

**Fig 7:**
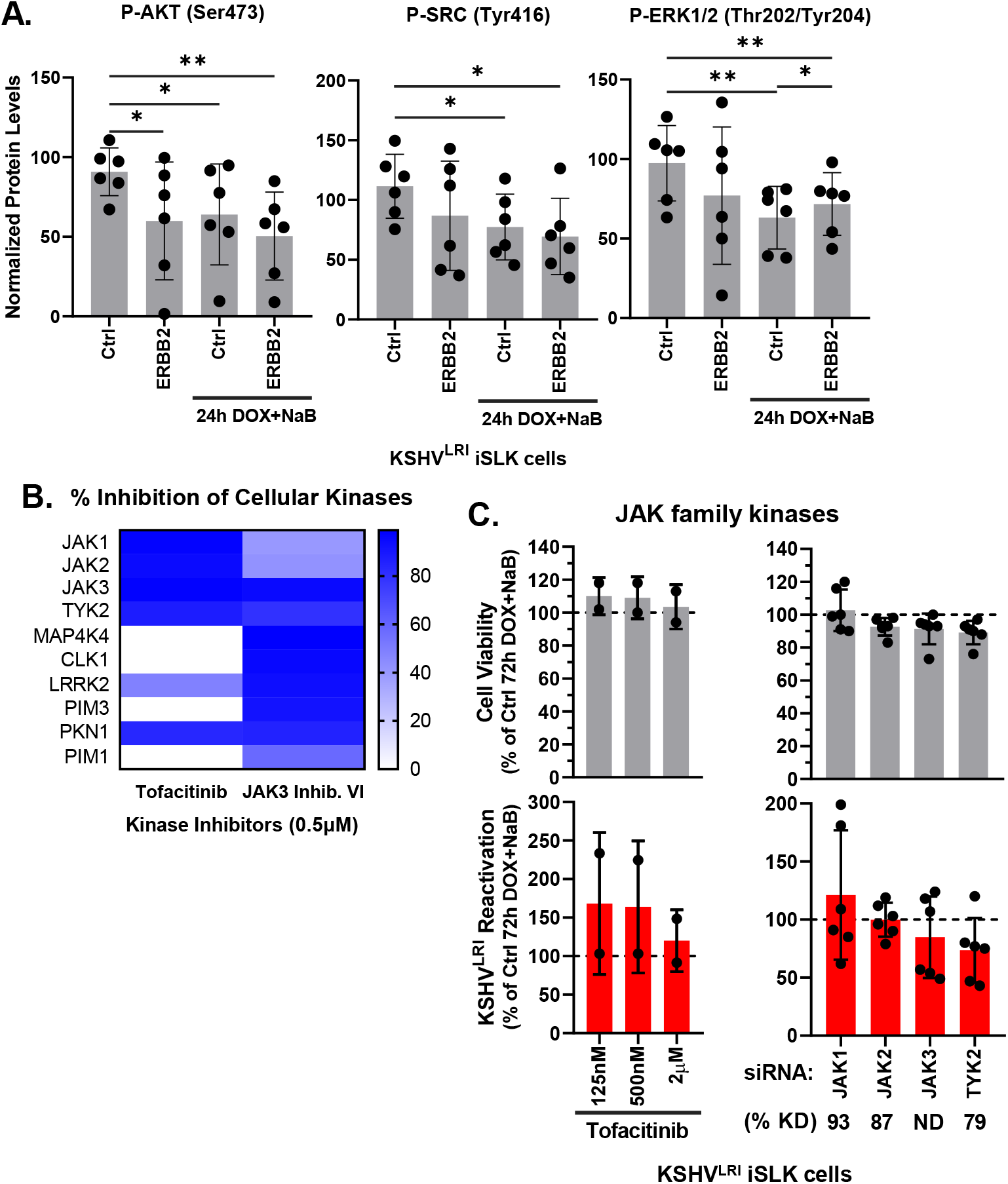
ERBB2 phosphorylation of intermediate kinases. **(A)** KSHV^LRI^ latently infected iSLK cells were transfected with siRNA control or siRNAs targeting ERBB2 and then 3-days later untreated or treated with DOX plus NaB for 24h. Cells were harvested, and protein lysates were analyzed using a RPPA for the phosphorylation of the indicated signaling intermediates downstream of ERBB2. **(B)** Kinase inhibition profiles for tofacitinib and JAK3 inhibitor VI from the Kinhibition database (https://kinhibition.fredhutch.org/). JAK3 VI inhibitor restricts JAK3, TYK2 and 15 other kinases. **(C)** Cell viability (grey bars) and KSHV reactivation (red bars) were measured for KSHV^LRI^ latently infected iSLK cells untreated or tofacitinib treated cells in combination with lytic inducing agents DOX plus NaB for 72h and for cells transfected with siRNAs targeting individual JAK family kinases and 3-days later treated with DOX plus NaB for 72h. Control DMSO or control siRNA transfected cells treated with DOX plus NaB (dotted black lines) were set to 100 and data for each condition was calculated as a percent of the control. Kinase knockdown efficiencies at 3-days following siRNA transfection were determined before addition of lytic inducing drugs and graphed in Fig S6. For each knockdown, the efficiencies were averaged and listed below the corresponding kinase target as % KD. P-values * ≤ 0.05 and ** ≤ 0.01.

The janus kinase family (JAKs) are also signaling intermediates downstream of the ERBBs. The JAK3 Inhibitor VI, predicted from the initial KiR screen to affect reactivation, was confirmed to regulate lytic reactivation (Fig 2D). To further investigate the role of JAK signaling, we tested another JAK inhibitor, tofacitinib that has greater specificity for the JAK proteins (Fig 7B) and found that it did not affect reactivation (Fig 7C). Since tofacitinib inhibits the catalytic activity of all JAK kinases while the JAK3 Inhibitor VI is more specific for JAK3 and TYK2 JAKs and several other unrelated kinases (Fig 7B), we wondered if different JAK family members may have counteracting effects on reactivation as we observed for the MKNK family protein kinases (Fig 4). To test this hypothesis, we used specific siRNAs to deplete JAK1, JAK2, JAK3, or TYK2 and performed assays under two lytic induction conditions. Under full lytic inducing conditions (DOX + NaB), none of these individual depletions impacted reactivation significantly (Fig 7C). We also conducted experiments in uninduced and DOX-only induced cells and observed that depletion of JAK1 increased reactivation 2.4-fold for the DOX condition as compared to the control (Fig S5C), suggesting that JAK1 is a pro-latency factor. Knockdown efficiency and specificity were shown for siRNAs targeting each JAK1, JAK2 and TYK2 using RT-qPCR (Fig S6A). JAK3 RNA abundance was below the level of detection for RT-qPCR. To assess the knockdown efficiency of the siRNAs targeting JAK3, we transiently overexpressed JAK3 by transfecting HeLa cells with a JAK3 expression plasmid and then transfected cells with targeting siRNAs followed by immunoblot analysis of JAK3 protein levels. Exogenously expressed JAK3 was almost completely depleted in this system (Fig S6B). These findings illustrate that the JAK1 protein kinase may have pro-latent activity which could be regulated like AKT, SRC and ERK1/2, by ERBB1:ERBB2 heterodimeric signaling or by another upstream receptor that is activated in response to RTA expression.

### ERBB2 phosphorylates CREB1, STAT1, and STAT3 transcription factors during KSHV reactivation

Finally, we tested the role of ERBB2 on the activation of proteins downstream of the selected signaling intermediates, including myristoylated alanine-rich C-kinase substrate (MARCKS), S6 ribosomal protein, and several transcription factors (Fig 6A). Phosphorylation of the MARKS substrate, which is downstream of PKC, was increased during induction of lytic replication (Fig S5D). ERBB2 knock down lessened the induction but not to a statistically significant extent. We detected only small and mostly insignificant effects of lytic induction and ERBB2 knock down on phosphorylation of S6, NFκB and β-catenin (Figs S5E and F). Despite these findings, we did observe a striking phenotype for three other transcription factors. Specifically, the cyclic AMP-responsive element-binding protein 1 (CREB1), signal transducer and activator of transcription 1 (STAT1), and STAT3 all exhibited a spike in activation under lytic inducing conditions as compared to latency and the effect was attenuated by depletion of ERBB2 in lytically induced cells, supporting a signaling function of ERBB2 to activate these transcription factors (Fig 8). In KSHV infected cells, phosphorylation of CREB1 Ser^133^ by MSK1/2 was reported to enhance KSHV lytic gene expression [18]. Thus, our results highlight the role of ERBB2 upstream of CREB1 activation to promote lytic gene expression, a previously undescribed connection during KSHV reactivation.

**Fig 8:**
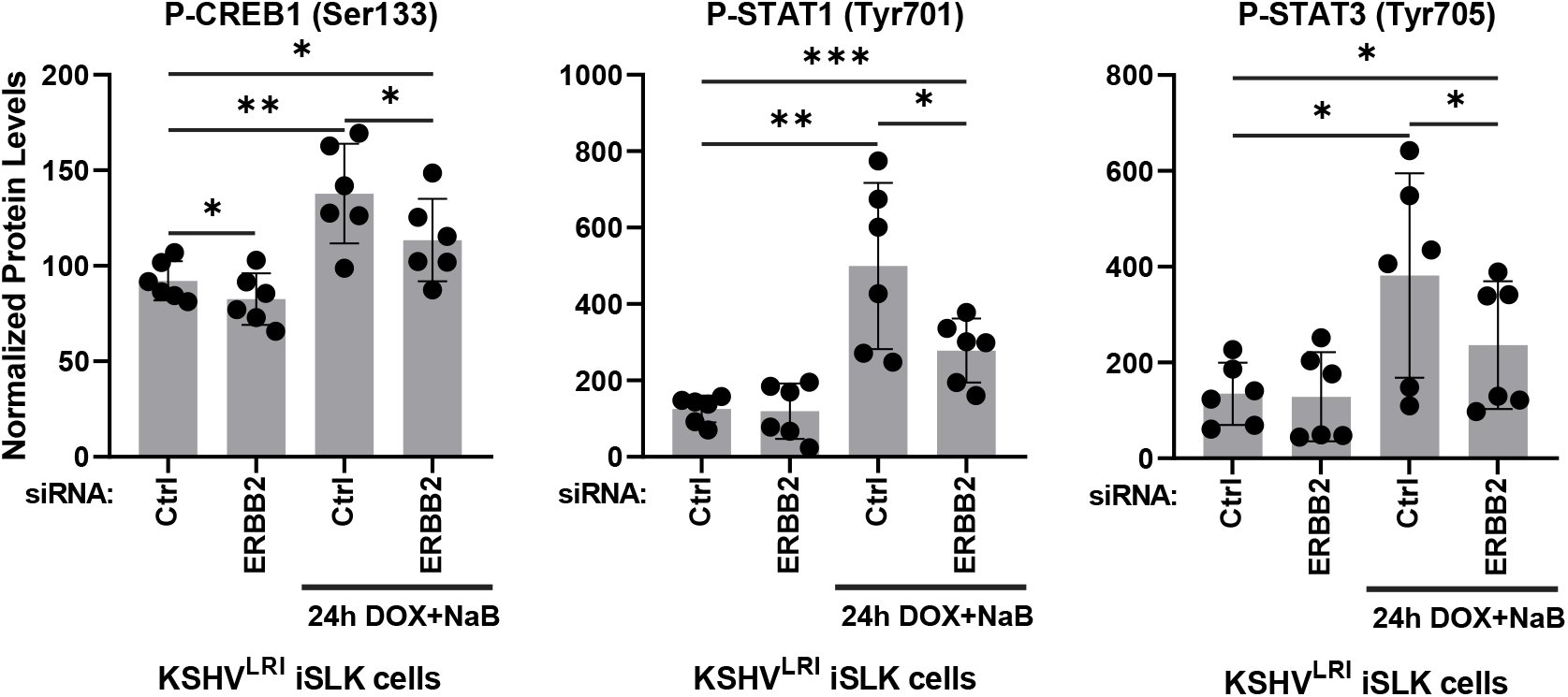
ERBB2-mediated signaling increases CREB1, STAT1 and STAT3 phosphorylation during lytic replication. KSHV^LRI^ latently infected iSLK cells were transfected with siRNA control or siRNAs targeting ERBB2 and then 3-days later untreated or treated with DOX plus NaB for 24h. Cells were harvested, and protein lysates were analyzed using a RPPA for transcription factor phosphorylation of CREB1 at Ser^133^, STAT1 at Tyr^701^, and STAT3 at Tyr^705^. P-values * ≤ 0.05, ** ≤ 0.01, and *** ≤ 0.001.

## Discussion

Protein kinases are known to regulate myriad host and viral processes. Like other viruses, KSHV relies on virus-encoded [44] and host-encoded kinases for optimal lytic replication [19, 21, 22, 45]. Some kinases regulate reactivation by altering LANA latent protein functions, lytic gene transcription and translation, and cell survival and proliferation [18-20, 23, 46-49]. Kinases with previously confirmed pro-lytic functions, including Pim-1/3 [19, 23], MSK1/2 [18], and RSK/ERK [20, 49, 50], appear to act by regulation of latency-associated nuclear antigen (LANA) phosphorylation, activation of CREB1 lytic gene transcription factor, and cell proliferation and survival, respectively. Kinases with pro-latency roles include CDK6 which interacts with KSHV v-cyclin and nucleophosmin to restrict lytic replication [48], and AMPKα1 which restricts lytic replication following primary infection through an unknown mechanism [46]. While the role of kinases as regulators of KSHV reactivation has been established, the previously published screens have limitations. For example, kinase overexpression screens can result in artificial kinase catalytic activity, localization, or protein interactions, surveys of protein phosphorylation can implicate activated signaling pathways with substrates shared by many kinases, and proteome analyses can inform on kinase abundance but not on catalytic activity or function [18, 23, 24]. Additionally, screening of KIs can result in many off-target effects due to the inherent broad activity of these drugs [25]. The KiR kinome screening approach that we employed is not without limitations but takes a unique approach as compared to these previously published screens to build on our understanding of kinases and KSHV reactivation. A key advantage of our method is the use of data characterizing the varying potencies of KIs for multiple targets to enable predictions of kinase inhibitors and specific kinases that regulate reactivation. As revealed, our report of this approach can provide new insights into kinase regulation of KSHV latency and reactivation.

Using the polypharmacology-based KiR screening method, we were able to predict and validate both kinase inhibitors and specific kinases that regulate KSHV latency and reactivation. Two of the top kinase inhibitors included lestaurtinib, a broadly acting tyrosine kinase inhibitor and K252a, a staurosporine analog that also has broad inhibitor potency. Neither of these drugs has been tested for effects on the KSHV latent-to-lytic replication switch, although lestaurtinib restricts multiple stages of adenovirus replication in cell culture [51] and K252a impedes EBV lytic replication [52]. None of the six initially validated kinases, ERBB4, MKNK2, ITK, TEC, DSTYK, and FRK (Fig. 3), had been specifically characterized previously as regulators of KSHV reactivation. The kinases that were predicted by the KiR screen but did not validate by siRNA knock down included several MAPKs (MAP3K8, MAP2K2, MAP4K4), CAMK2G, and LRRK2, all of which have functional paralogs. It is possible that redundancies in the signaling pathways would require knocking down more than one of these kinases to reveal a reactivation phenotype. For example, CAMK2G forms complexes with the other CAMK2 paralogs to generate a 12-14 subunit holoenzyme [53], which might retain function if only one member is depleted. Additionally, LRRK2 may be essential for cell survival as suggested by poor knockdown efficiencies and reduced cell viability in some experiments. Unlike the other kinases that did not validate from the screen, the PBK kinase does not have a paralog, but it has multifunctional roles in regulating cell cycle progression. Therefore, knockdown of PBK may dysregulate opposing signaling pathways to disguise PBK functions during lytic replication. Lastly, the CLK1 kinase had poor knockdown efficiency so we could not evaluate its role in reactivation. While siRNA targeting CLK1 mRNA was published to knockdown both pre- and mature CLK1 mRNA relatively well, restoration of depleted CLK1 RNA occurs rapidly under cell stress conditions [54] which may in part explain the poor knockdown in KSHV infected cells. Despite these limitations, siRNA-mediated knockdown of six kinases validated these as newly described regulators of KSHV reactivation.

The two validated screen hits with the strongest impacts on reactivation, ERBB4 and MKNK2, are from kinase families with members that share high sequence homology and functional overlap. ERBB family members have well-characterized roles in cancer, cell proliferation, and cell survival, and they can form heterodimers to regulate the signaling pathways for these biological processes. MKNK1, like MKNK2, can phosphorylate the cap-binding protein eIF4E to regulate translation [55]. MKNK2 has also been shown to phosphorylate KSHV LANA latent protein in *in vitro* kinase assays [56]. To determine if these closely related kinases have pro-lytic functions like ERBB4 and MKNK2, we tested their effect on KSHV reactivation. Indeed, knocking down either ERBB2 (also called HER2 or *neu*) or ERBB3 inhibited reactivation, even somewhat more strongly than ERBB4 (Fig 4A). In contrast, knocking down ERBB1 (Figs 5A and 5B) or MKNK1 (Fig 4B) promoted certain stages of lytic replication, suggesting that they positively contribute to latency maintenance. These studies illustrate that kinase paralogs with some shared signaling pathways, nonetheless, can have counteracting roles in regulating the KSHV latent-to-lytic replication switch.

We selected ERBB2 for further investigation because it plays a unique function in ERBB complexes as a dominant binding partner that drives prolonged signaling [36] and, in some cancers, behaves as an oncogene [57]. We reasoned that study of ERBB2 signaling may provide insight into how latency and/or lytic replication mediates KSHV-dependent oncogenesis. In latently infected iSLK cells, ERBB1 and ERBB2 are expressed while ERBB3 and ERBB4 are detectable only after treatment of cells with lytic inducing agents (Table S4, [34]). These published data suggest that during latency, only ERBB1 is available for binding to ERBB2. Consistent with this possibility, ERBB2 knockdown reduced phosphorylation of ERBB1 at Tyr^1173^ during latency, but not in cells with activated lytic replication (Fig 6B). These data, along with the finding that ERBB1 restricts lytic gene expression (Fig 5), support a model in which ERBB1 restriction of lytic gene expression occurs via ERBB1:ERBB2 heterodimer signaling during latent infection but during lytic replication, this interaction is disrupted (Fig. 9). Intriguingly, this dissociation event coincides with the expression of ERBB3 and ERBB4 which could facilitate ERBB2 partner switching, especially because ERBB2 has a stronger affinity for ERBB3 binding as compared to the other ERBBs [36].

**Fig 9.**
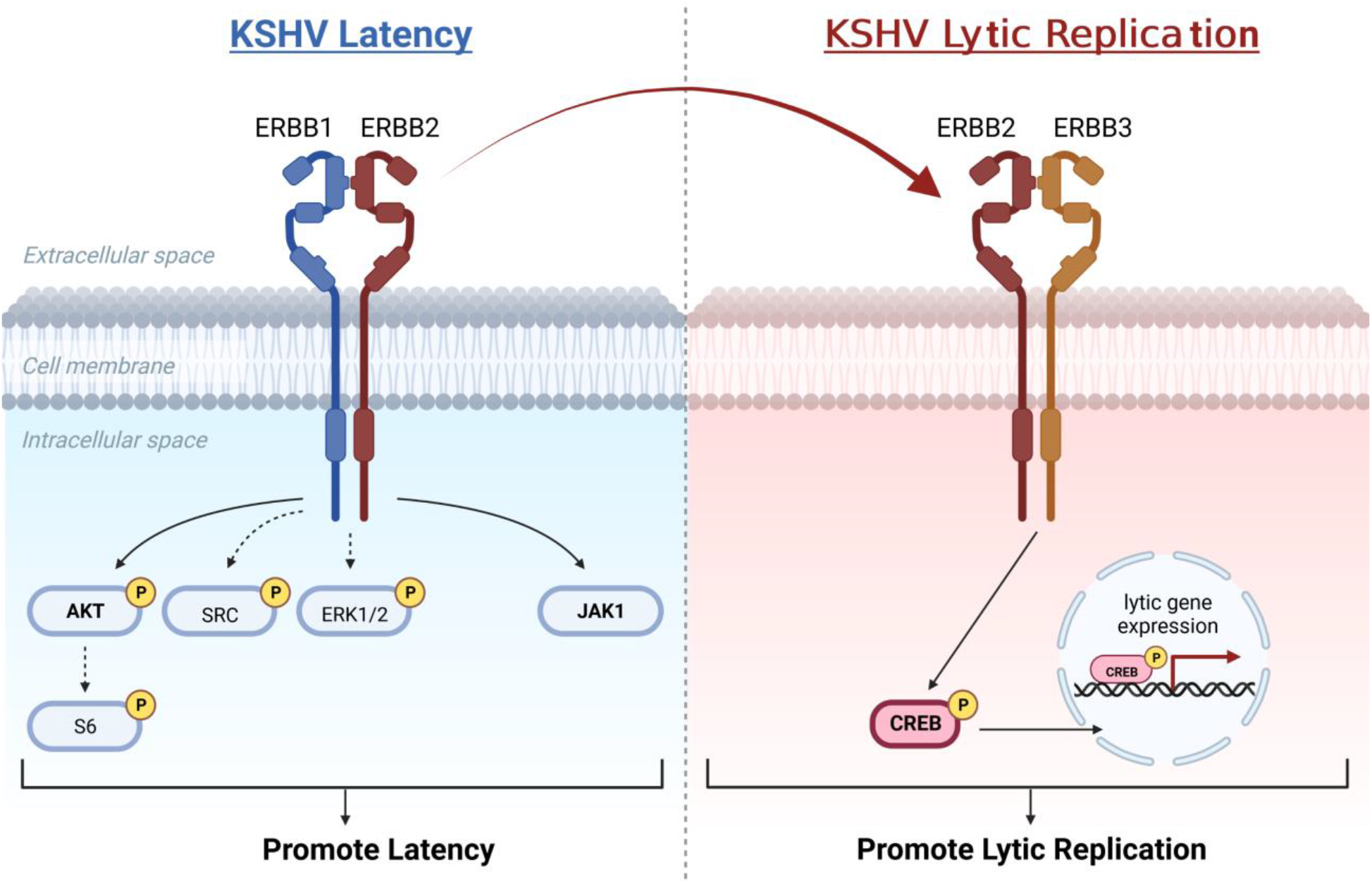
Model of ERBB family kinase roles in regulating KSHV latent-to-lytic replication switch. ERBB1:ERBB2 heterodimer signaling activates AKT and this trend applies to S6, SRC and ERK1/2 signaling intermediates to promote the latent state. JAK1 in some contexts also promotes latency and may function downstream of the ERBB1:ERBB2 heterodimer. This signaling is repressed during lytic replication as ERBB2 forms a heterodimer with newly expressed ERBB3, a switch facilitated by high affinity binding of ERBB2 to ERBB3, and ERBB2-dependent signaling activates lytic gene transcription factor, CREB1.

To determine which of the many signaling pathways regulated by the ERBB kinases are ERBB2-dependent during latency and lytic replication, we assayed for the phosphorylation of key residues that are indicative of activation for a subset of signaling factors. The selected proteins included plasma membrane proteins involved in crosstalk with ERBB kinases, downstream signaling intermediates of ERBB kinases, and downstream substrates of the signaling intermediates (Fig 6A). Activation of plasma membrane receptors may contribute to the ERBB2 pro-lytic mechanism via receptor crosstalk pathways. For example, the mesenchymal epithelial transition (MET) oncogene can activate ERBB1 in some cancer cells and in others it is activated by ERBB1:ERBB2 heterodimers [40, 42]. Also, signaling by the platelet derived growth factor receptor beta (PDGFRβ) overlaps with ERBB signaling intermediates [43]. Our data show that MET signaling decreases with reactivation through an ERBB2-independent mechanism, while PDGFRβ phosphorylation was unchanged under all tested conditions (Figs 6C and S5A). These findings do not support a role for MET or PDGFRβ as downstream factors of ERBB1:ERBB2 signaling.

From the investigation of signaling intermediates, we found further evidence consistent with ERBB1:ERBB2 signaling during latency. In cells containing latent virus, AKT activation was reduced by ERBB2 depletion and other intracellular kinases, SRC and ERK1/2, showed similar although not statistically significant trends (Fig. 7A). Contrary to the latent state, during lytic replication ERBB2 does not appear to activate these signaling intermediates. We also assayed for the role of JAK family kinases as these kinases are intermediates of ERBB signaling cascades and the JAK3 Inhibitor VI restricted reactivation (Fig 2D). The individual JAK family kinases did not have pro-latent or pro-lytic phenotypes under DOX-induced RTA plus NaB conditions (Fig 7C), but we did observe a moderate pro-latent phenotype for JAK1 under DOX-induced RTA alone conditions (Fig S5C). Our investigation of ERBB2 sigaling revealed that ERBB1:ERBB2 heterodimer activates AKT and to a lessor extent SRC and ERK1/2 during latency. JAK1 may also be activated by these receptors or another factor to promote latency. But none of these signaling intermediates likely contribute to the ERBB2-driven promotion of lytic replication.

The last category of factors that we tested are downstream substrates of the selected signaling intermediates, although we recognize the caveat that other factors may regulate these substrates. We found that ERBB2 was required to fully activate CREB1 (Fig 8), a transcription factor that promotes KSHV lytic gene expression [18]. ERBB2 also activated STAT1 and STAT3 during lytic replication (Fig 8), but only pro-latency roles for STAT1 and STAT3 have been described [58]. Intrestingly, in some cancer cells, STAT1 can negatively regulate ERBB2/Neu-dependent transformation [59]. If this is the case in KSHV infected cells, STAT1 may provide a negative feedback loop to dampen ERBB2 signaling [60]. Future investigation of the ERBB2-STAT1 signaling axis may inform on whether or not ERBB2 regulates cellular transformation in KSHV infected cells. Our working model of ERBB siganling supported by these studies suggests that ERBB1:ERBB2 heterodimers form and activate AKT and likely other downstream intermediates to promote KSHV latency. The induction of lytic replication activates ERBB3 expression providing a new, high-affinity binding target for ERBB2 that competes with ERBB1 for ERBB2 binding. We propose that ERBB3 presence in part activates KSHV replicative switch by causing ERBB2 dissociation from ERBB1 and the formation of ERBB2:ERBB3 heterodimers that activate CREB1 lytic gene transcription factor, ultimately promoting KSHV lytic replicative cycle (Fig 9).

While our signaling cascade analyses only revealed activation of CREB1 downstream of ERBB2 during lytic replication, important signaling intermediates of ERBB2-dependent signaling may include the other KiR screen hits. The ubiquitously expressed MKNKs are activated by p38 (or ERK1/2), which are part of the MAPK signaling pathways downstream of the ERBBs. Thus, testing the role of ERBB2 upstream of these other kinases will be of interest. The expression of another pro-lytic kinase validated from our screen, ITK, is induced following KSHV reactivation in iSLK cells (Table S4, [34]). This kinase and other TEC family kinases can interact with ERBB3 cytoplasmic tail following ERBB2-mediated phosphorylation [61, 62] suggesting that TEC family kinases may function as signaling intermediates during the transition between latency and lytic replication. Continued dissection of the roles of kinases during KSHV lytic replication will provide insight into the overlapping or parallel pathways at work to coordinate this replicative switch of KSHV.

Together, these experiments confirm the utility of a polypharmacology-based kinome screen to study KSHV reactivation regulators. The iSLK system provided a convenient cell line for these proof-of-principle studies with relatively high levels of induced-lytic replication, which is not achievable in most KSHV latent cell culture systems, though conducting this screening approach in other cell types and conditions is of interest. The translational potential of this research is most evident with the new connection identified between KSHV latent-to-lytic replication switch and ERBB signaling. Viruses encoding or overexpressing ERBB1 agonists demonstrate the role of ERBB1 signaling during viral life cycles including virus-mediated tumorigenesis [63–65]. Also, the ERBBs are well-studied regulators of tumorigenesis and have several targeted, FDA-approved therapies in use for some cancers [16, 57, 66, 67]. Our investigation of kinase regulators revealed for the first-time counteracting roles of the ERBBs to coordinate the critical KSHV latent-to-lytic replication switch. Continued mechanistic probing of these factors will enhance our understanding of the intricacies of this viral switch and application to other cell types and systems may inform on the therapeutic potential of targeting these kinases to affect KSHV-driven diseases.

## Materials and Methods

### Cell culture

The renal carcinoma cell line (SLK) and doxycycline inducible RTA SLK (iSLK) cell line were a kind gift from Jae Jung (Cleveland Clinic) and Rolf Renne (University of Florida). The SLK, iSLK, 293T, and HeLa cell lines were maintained at 37°C and 5% CO_2_ atmosphere in Dulbecco’s modified Eagle’s medium (DMEM) supplemented with 10% NuSerum (Corning #355500) and 1% penicillin/streptomycin (Gibco #15140122). For iSLK cells, media was supplemented with 1 µg/ml puromycin (ThermoFisherScientific #BP2956100) and 250 µg/ml G418 (Sigma #A1720). After infection of iSLK cells, KSHV BAC16 or KSHV^LRI^ episomes were maintained by addition of 1,000 µg/ml hygromycin B (Invitrogen #10687010) to the medium. All cell lines were confirmed to be mycoplasma negative using the MycoProbe kit (R&D Systems #CUL001B).

### Generation of KSHV^LRI^ recombinant genome

The KSHV^LRI^ recombinant genome was generated by recombining a dual lytic replication indicator cassette into the KSHV BAC16 genome. The KSHV BAC16 genome was generously provided by Jae Jung in the GS1783 *E. coli* strain [68]. The dual lytic replication indicator cassette was generated by PCR amplification of the KSHV *PAN* promoter (PrPAN) using BAC16 DNA as the template and primers 2580 and 2581 (Table S5) and inserting the amplicon into the pcDNA3.1 V5-His-TOPO vector. The lytic replication indicators included a streptavidin-binding peptide fused to a truncated low nerve growth factor receptor (SBP-ΔLNGFR) and a nuclear-localized mCherry (mCherry-nls) separated by a P2A sequence. The SBP-ΔLNGFR was amplified from pJB-2045_CMV_SBPΔLNGFR (a gift from Jesse Bloom, Fred Hutchinson Cancer Center, [69] with primers 2585 and 2586. The P2A-mCherry-nls was amplified from pEH_mCherry-NLS-TagRFP plasmid (a gift from by Emily Hatch, Fred Hutchinson Cancer Center, [70]) with primers 2600 and 2601. The PrPAN, SBP-ΔLNGFR, P2A-mCherry-nls DNA fragments were cloned into the pcDNA3.1 V5-His-TOPO vector at HindIII, KpnI/BAMHI, and BAMHI/NotI sites respectively.

To seamlessly introduce the lytic indicators into the KSHV BAC16, an additional I-SceI sequence [71] was cloned into the BamHI site between the SBP-ΔLNGFR and P2A sequences. Upstream of the I-SceI cleavage site, a 50bp DNA segment identical to the P2A sequence was added to the forward primer. The I-SceI-KanR sequence was amplified from pEPKan-S2 (provided by Greg Smith (Northwestern University, [71] with primers 2610 and 2611, where the forward primer contained the 50bp overlapping sequence and both primers contained BAMHI sites on the outer flanks. This DNA segment was cloned into the BAMHI site of the pcDNA3.1 PrPAN-SBP-ΔLNGFR-P2A-mCherry-nls intermediate to make the pEQ1766 plasmid. Next, the PrPAN-SBP-ΔLNGFR-P2A-50bp-I-SceI-KanR-mCherry-nls gene cassette was amplified from pEQ1766 using primers 2617 and 2618. This cassette was inserted into the KSHV BAC16 genome containing GS1783 *E.coli* by seamless recombineering as described in [71, 72].

The sequences of the KSHV^LRI^ BAC and of BAC16 genomes were verified by Illumina deep sequencing. Briefly, 100 ng purified genomic DNA from a BAC16 or KSHV^LRI^ clone were used to generate libraries using the KAPA HyperPlus kit and sequenced using an Illumina MiSeq. Reads were trimmed using Trimmomatic v0.39 and mapped to the human herpesvirus 8 strain JSC-1 clone BAC16 reference genome GQ994935.1 using Geneious read mapper [73, 74]). Sequencing reads were deposited in NCBI BioProject PRJNA884721.

### Generation of stable KSHV^LRI^ latently infected iSLK cells

KSHV latently infected iSLK cell lines were generated by first transfecting 293T cells with purified BAC16 or KSHV^LRI^ DNA and subsequent co-culture with uninfected iSLK cells, as described by Jain et al. [75]. After selecting the transfected 293T cells with 100 µg/ml hygromycin, the virus was reactivated by adding 20 nM phorbol 12-myristate 13-acetate and 1 mM valproic acid to the medium. After 48-72h, the virus containing medium plus 8 µg/ml polybrene (Sigma #H9268) and the infected 293T cells were co-cultured with iSLK cells. Several days later, the co-culture media was changed to media supplemented with 500 µg/ml hygromycin, 1 µg/ml puromycin and 250 µg/ml G418 to select for KSHV latently infected iSLK cells. These cells were further selected with 1,000 µg/ml hygromycin and frozen in liquid nitrogen after two or three passages. Cells used for experiments were passaged less than 10 times.

### KSHV reactivation mCherry fluorescence assay

KSHV^LRI^ latently infected iSLK cells with no prior treatment or those treated with siRNAs at 2-days post transfection were seeded into 96-well plates at 2.5×10^4^ cells per well. The next day, cells were untreated, treated with 1 µg/ml DOX or treated with DOX and 1mM NaB for controls, or at the same time treated with kinase inhibitors for drug experiments. At 72 h post treatment, mCherry fluorescence object count per image was quantified using an Incucyte Imaging System (Sartorius). Cell viability was determined by percent cell confluence from phase images or GFP as measured by the Incucyte as an average per well. For siRNA treated cells, data in which the cell viabilities were > 1.5 standard deviation (>31% reduction as compared to siCtrl cells under DOX and NaB conditions) were removed from analysis. One replicate was removed for MKNK1, MKNK2, and LRRK2 siRNA mediated depletion experiments because of poor cell viability.

### Kinase inhibitor treatment

KSHV^LRI^ latently infected iSLK cells were seeded at 2.5×10^4^ cells per well into 96-well plates. The next day, the medium was replaced with medium containing the KI alone, KI plus 1µg/ml DOX, or KI plus DOX and 1mM NaB. Control wells included medium with the vehicle control (DMSO at 0.2% or less) in place of the KI. Each treatment was conducted with 3 technical replicates. The initial training set of KIs described in Gujral et al [31] were tested at 2µM, 500nM, 125nM and 31nM concentrations. All 29 proprietary KIs for the initial training set (Gujral 2014) were tested once and 12 of these were tested twice and the two experiments were averaged. The validation set of KIs and tofacitinib (Table S1) were tested at 2µM, 500nM, and 125nM in two or three separate experiments. All small molecule KIs for the initial training set were constituted in DMSO at 1mM stock solutions (0.2% DMSO for 2µM concentrations). For the validation set, KIs and tofacitinib were constituted in DMSO at 10mM or 4mM stock solutions (0.02% or 0.05% DMSO for 2µM concentrations).

### Kinase inhibitor Regularization (KiR) modeling

KiR models for KSHV reactivation were generated as previously described [31, 33]). A set of 29 inhibitors were tested on KSHV^LRI^ latently infected iSLK cells as described above, with the end result being a single response for each drug that represents the change in KSHV reactivation (as % DMSO control) at the profiled dose of the inhibitor. The kinase inhibition profiles of each inhibitor and the quantitative responses to those inhibitors were used as the explanatory and response variables, respectively, for elastic net regularized multiple linear regression models[76]. Custom R scripts (available at https://github.com/FredHutch/KiRNet-Public) employing the glmnet package were used to generate the final models. [77] Leave-one-out cross-validation (LOOCV) was used to select the optimal value for the penalty scaling factor λ. Models were computed for 11 evenly spaced values of α (the relative weighting between LASSO and Ridge regularization) ranging from 0 to 1.0 inclusive. Kinases with positive coefficients in at least one of these models (with the exception of α = 0, which always has non-zero coefficients for every kinase) were considered hits.

### siRNA or plasmid transfections

KSHV^LRI^ latently infected iSLK cells were seeded at 4×10^5^ cells/well into a 6-well plate. The next day, cells were transfected with 100nM non-targeting control or individual kinase targeting On-TARGETplus SMARTPool siRNAs (horizon; Table S6) including four unique target sequences and 5µl RNAiMAX (ThermoFisher #13778150) in Opti-MEM (ThermoFisher #31985088) per well following manufacturer’s instructions. For transfection with the JAK3 expressing plasmid, HeLa cells were seeded into a 6-well plate at 5×10^5^ cells/well. The next day, cells were transfected with siRNAs as described above and 2 days following cell seeding, cells were transfected with 1.5µg pcDNA3.1-JAK3 and 3µl lipofectamine 2000 (ThermoFisher #11668027) per well in Opti-MEM following manufacturer’s recommendations.

### Reverse transcription qPCR

Knockdown efficiencies of targeting siRNAs were evaluated by reverse transcription qPCR 3-days after siRNA transfection. For relative quantification of lytic gene expression, cells transfected with siRNAs were treated with 1 µg/ml DOX plus 1 mM NaB at 3-days post transfection and harvested at 24h following lytic induction for *mCherry* and *ORF10* and at 48h for *K8.1.* Total RNA was extracted from cells following the RNAeasy Mini Kit (QIAGEN #74104) protocol and cDNA was synthesized from RNA samples according to the High-capacity RNA-to-cDNA (AppliedBiosystems #4387406) protocol. Quantitative PCR of cDNA samples were carried out using Power Sybr Green Master Mix (ThermoFisher #4367659) on a Bio-Rad CFX384 Real Time System C1000 Touch Thermal Cycler using primers listed in table 5. Relative mRNA levels were normalized to tubulin control and calculated using the ΔΔCt method for experimental conditions as compared to control conditions.

### Reverse-phase protein arrays (RPPA)

Control or siRNA treated KSHV^LRI^ iSLK cells were harvested 4-days following siRNA transfection and 24h following KSHV lytic induction with 1 µg/ml DOX and 1 mM NaB for protein lysate microarray analysis. Sample preparations and protein array analyses were performed as detailed in Luckert et al. [78]. In brief, cells were rinsed then lysed in 50 mM Tris-HCl, 2% sodium dodecyl sulfate, 5% glycerol, 5 mM ethylenediaminetetraacetate, and 1 mM sodium fluoride, 1X Complete Protease Inhibitor Cocktail (1 tablet per 10 ml, Roche), 1X Pierce protease plus phosphatase inhibitor tablet (Thermo Scientific #A32959), 10 mM β-glycerol phosphate, 1 mM PMSF, 1 mM sodium orthovanadate, and 1 mM dithiothreitol. After filtering through a 0.2-µm filter plate, the lysates were printed onto a nitrocellulose-coated slide using Aushon 2470 arrayer. Primary antibodies listed in table S7 were diluted 1:100 and incubated with slide for 24h on an orbital shaker at 4°C. IRDye secondary antibodies listed in table 7 were diluted to 1:1000 and incubated with slide for 1h, shaking at room temperature. The microarray slides were scanned in 680-nm and 800-nm channels with an Odyssey imager. Protein quantitation for siCtrl, uninduced KSHV^LRI^ iSLK samples were set to 100 and samples were normalized to these controls. Six independent siRNA transfections were completed for the six RPPA experiments.

### Statistics

The statistical analyses for these data were completed using Graphpad Prism 8.0.1 software. Experimental conditions were compared to control conditions using 2-way ANOVA in GraphPad Prism 8.0.1 or unpaired t test in excel (* ≤ 0.05; ** ≤ 0.01; *** ≤ 0.001).

### JAK3 plasmid clone

The *JAK3* gene was amplified using primers 2636 and 2648 from the pDONR223-JAK3, a gift from William Hahn & David Root (Addgene plasmid # 23944; http://n2t.net/addgene:23944; RRID: Addgene_23944). The amplicon was cloned into the pcDNA3.1 V5-His-TOPO vector to generate the pEQ1797 plasmid. The sequence of the JAK3 insert in the resulting plasmid was verified by Sanger sequencing in the Fred Hutch Genomics Core.

### Immunoblot assays

Lysates of uninfected, KSHV^LRI^ latently infected, or infected and 3-day lytically induced iSLK cells were separated on 8 or 10% polyacrylamide gels. For ERBB2 and paired actin blots, KSHV^LRI^ latently infected iSLK cells were harvested 2 days after siRNA transfection. For the JAK3 and paired actin control blots, HeLa cells were harvested 3 days after siRNA transfection and 2 days after pcDNA3.1-JAK3 transfection. For all gels, the protein was transferred onto a polyvinylidene difluoride (PVDF) membrane (Millipore), and proteins were detected by probing with specific antibodies (Table S7) using the Western Star chemiluminescent detection system (Applied Biosystems) according to the manufacture’s recommendations.

### Immunofluorescence assay

For LANA IFA, uninfected or KSHV^LRI^ latently infected iSLK cells were seeded into an 8-well plastic chamber slide (ThermoFisherScientific #177445) at 2×10^4^ cells/well. The next day, cells were fixed with 4% paraformaldehyde (Electron Microscopy Sciences #15710) in 1X PBS for 15m. Cells were permeabilized and blocked simultaneously for 20m with 1% Triton X-100 (Sigma #X100), 0.5% Tween20 (ThermoFisherScientific #J20605.AP), 3% BSA (Sigma #A7906) and 1X PBS (Gibco #14200075) solution. Nuclei were stained using DAPI containing mounting medium (VECTASHIELD #H-1200-10). For SBP-ΔLNGFR IFA, KSHV^LRI^ infected iSLK cells were seeded into a 24-well plate at 8.0×10^4^ cells/well. The next day, the cells were treated with DOX plus NaB. 5-days following treatment, cells were fixed as stated above, blocked with 3% BSA in 1X PBS, and incubated with streptavidin Fluor 680 conjugate (Invitrogen #S21378). Cells were imaged on a Leica Microsystems DM IL LED Fluorescent microscope with Leica Application Suite V4.12 software.

### Kinase activity profiles for JAK inhibitors

The kinase activity profiles for tofacitinib and JAK3 inhibitor VI (Fig 7B) were taken from the publicly available Kinhibition website (https://kinhibition.fredhutch.org/). Tofacitinib data indicates specific restriction of all JAK kinases and two other kinases, LRRK2 and PKN1. The JAK3 VI inhibitor restricts JAK3, TYK2 and 15 other kinases including Pim-1 and Pim-3 pro-lytic kinases and two kinases predicted from the kinome screen, CLK1 and MAP4K4. A heatmap of kinase activities during treatment with 500nM of either tofacitinib or JAK3 inhibitor VI was generated in GraphPad Prism 8.0.1.

## Acknowledgments

This work was supported by grants from the Fred Hutchinson Cancer Center Pathogen-associated Malignancies Integrated Research Center, from the Fred Hutchinson Cancer Center Human Biology Division, from the National Institutes of Health grant R01AI45945 (to A.P.G.), R01CA189986, R01CA217788, and R21CA240479 (to M.L.), from the National Science Foundation grant 2047289 (to T.S.G.), and by the Fred Hutch/University of Washington Cancer Consortium (P30 CA015704). The content is solely our responsibility and does not necessarily represent the official views of the National Institutes of Health. The funders had no role in study design, data collection and analysis, decision to publish, or preparation of the manuscript.

The authors acknowledge the help of Drs. Alex Greninger and Nicole A. Lieberman from the University of Washington for help sequencing the KSHV^LRI^ recombinant.

## Figure Legends

**Fig S1:**
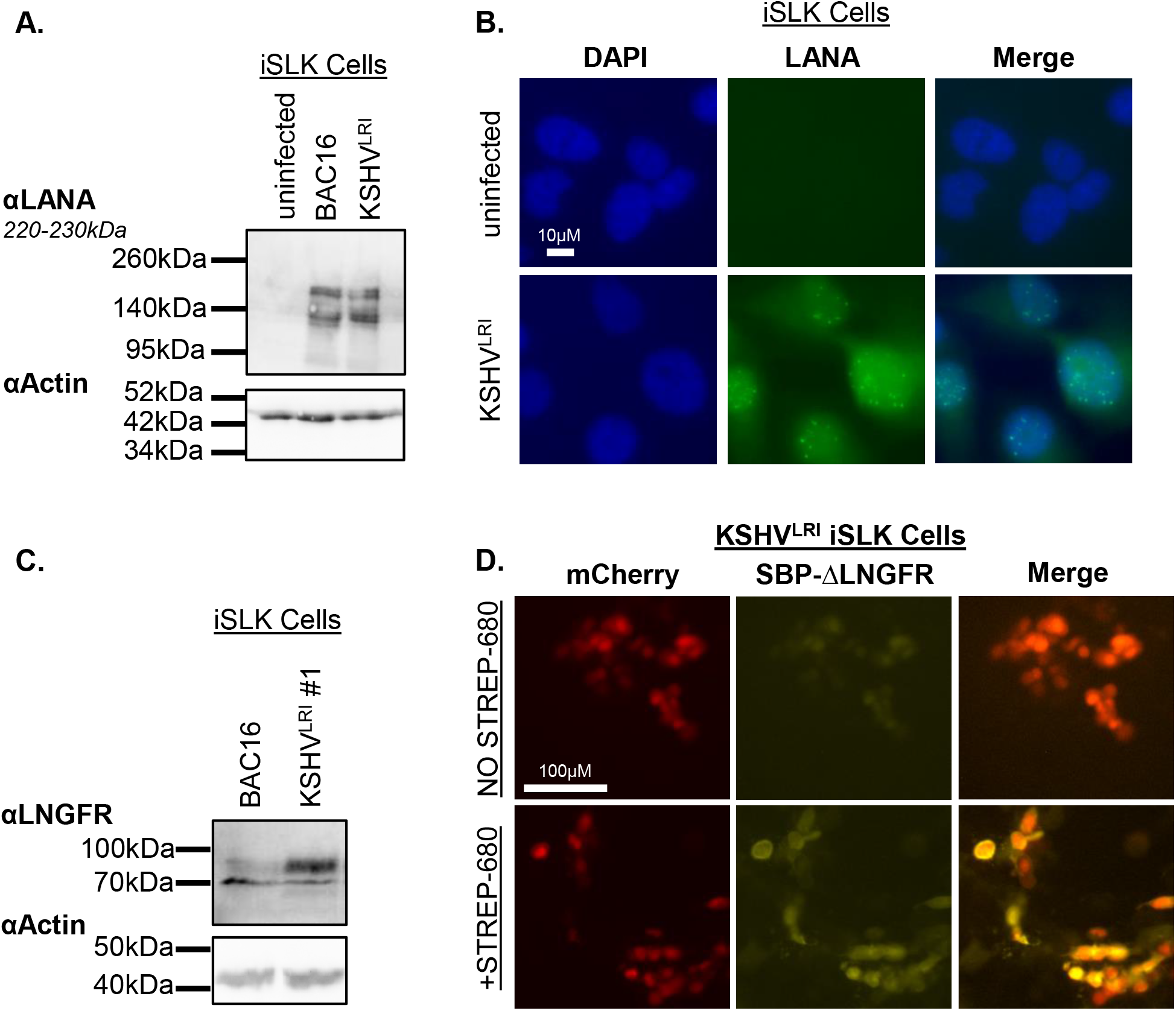
Expression of LANA and the lytic replication indicator SBP-ΔLNGFR from KSHV^LRI^. Uninfected iSLK cells or KSHV BAC16 or KSHV^LRI^ latently infected iSLK cells were **(A)** lysed and subjected to LANA immunoblotting or **(B)** analyzed by immunofluorescence for LANA puncta representing individual KSHV episomes. **(C)** KSHV BAC16 or KSHV^LRI^ latently infected iSLK cells were treated with 1 µg/ml DOX plus 1 mM NaB and incubated for 3-days before harvesting cells for immunoblot analysis of SBP-ΔLNGFR protein levels or **(D)** fixed and incubated with streptavidin-680 for imaging of SBP-ΔLNGFR on the plasma membrane of un-permeabilized cells.

**Fig S2:**
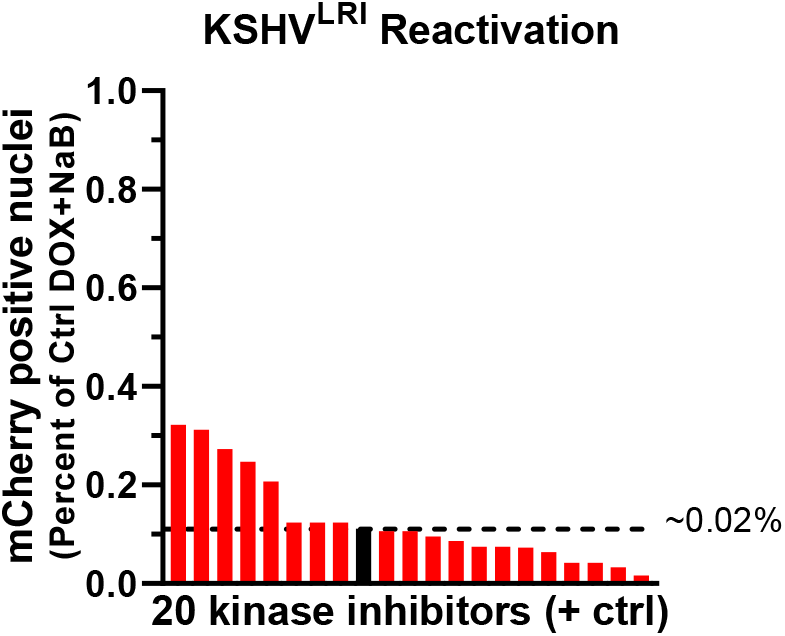
Polypharmacology-based kinome screen in the absence of KSHV lytic inducing agents. KSHV^LRI^ reactivation phenotypes were obtained from 20 of the 29 pre-selected kinase inhibitors that did not cause cellular toxicity. KSHV reactivation for control (black bar and dotted black line) and kinase inhibitor treatment (red bars) were calculated as a percent of DOX plus NaB treated cells set to 100 from data in Fig 2C. In this graph, 1.0 represents ∼0.2% total cells and the dotted line represents spontaneous reactivation, ∼0.02% total cells.

**Fig S3:**
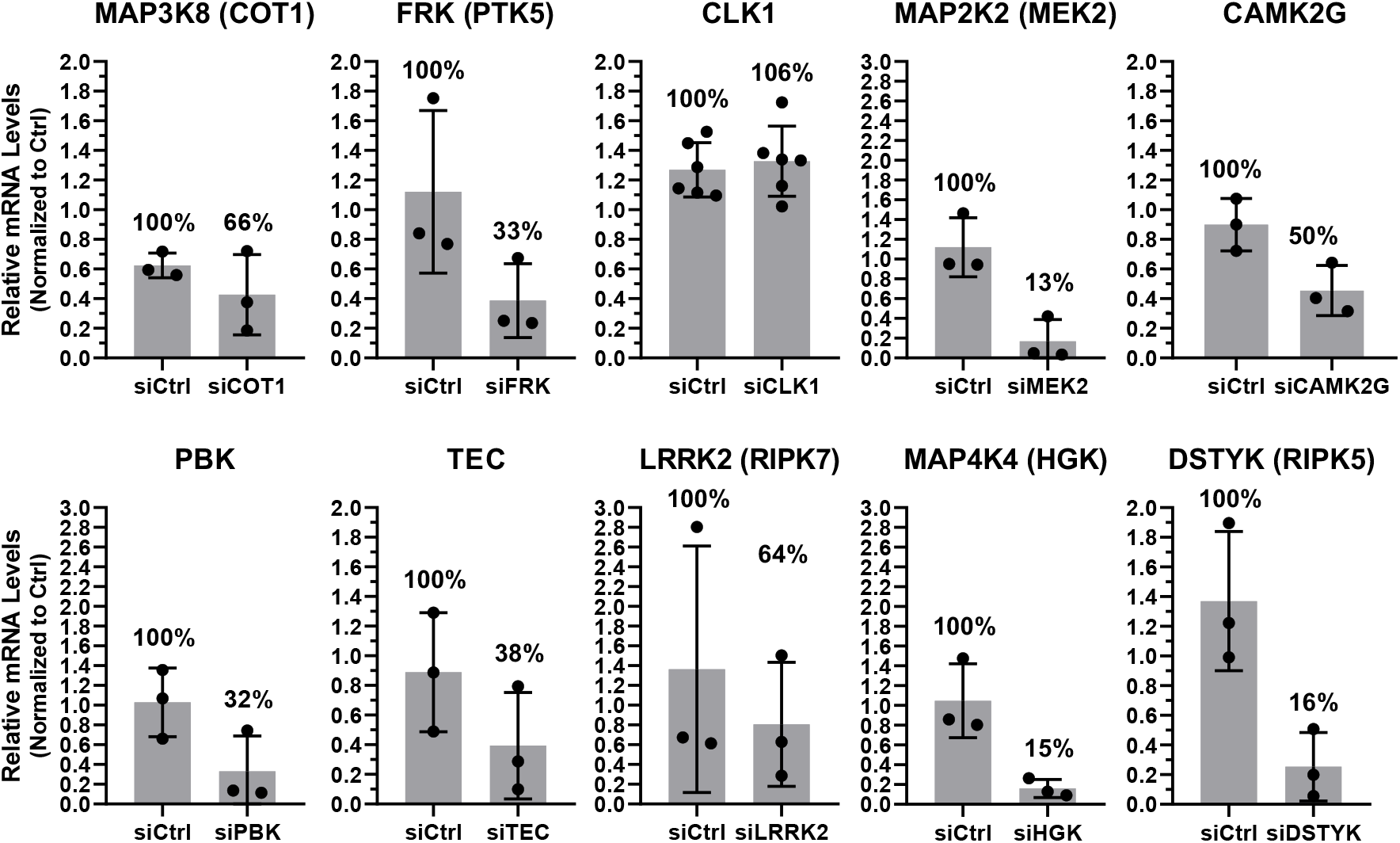
Kinase knockdown efficiencies for kinases validated from screen. Knockdown efficiencies for siRNAs targeting specific cellular kinases were evaluated in KSHV^LRI^ latently infected iSLK cells using RT-qPCR from total RNA harvested at 3-days post transfection with siRNAs.

**Fig S4:**
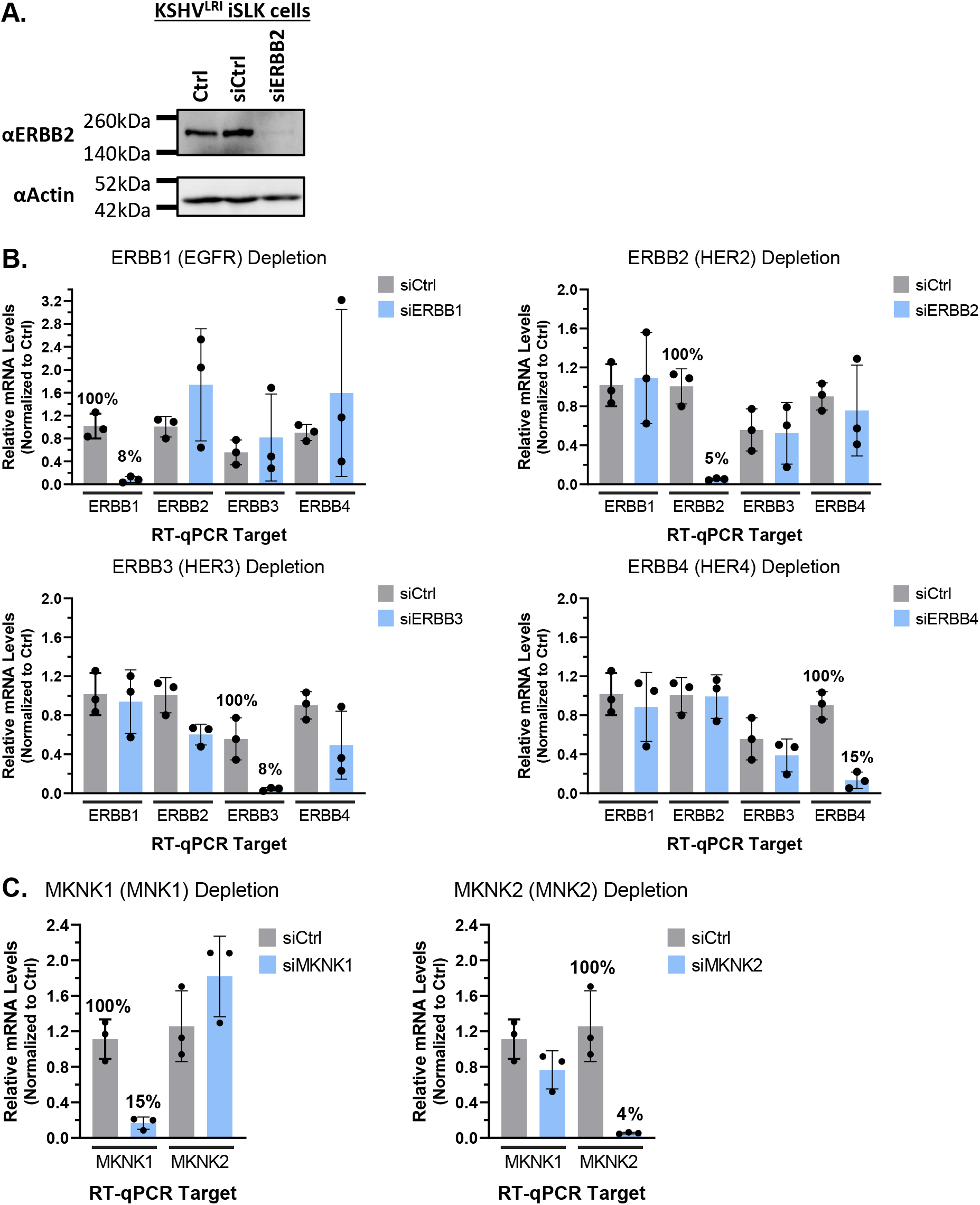
Knockdown efficiencies and specificity for ERBB and MKNK family members. **(A)** Knockdown efficiency of ERBB2 targeting siRNA was evaluated by immunoblot for ERBB2 protein at 2 days following siRNA transfection of KSHV^LRI^ latently infected iSLK cells. Knockdown specificity for siRNAs targeting **(B)** ERBB or **(C)** MKNK family kinases were evaluated in KSHV^LRI^ latently infected iSLK cells using RT-qPCR from total RNA harvested at 3-days post transfection with siRNAs.

**Fig S5:**
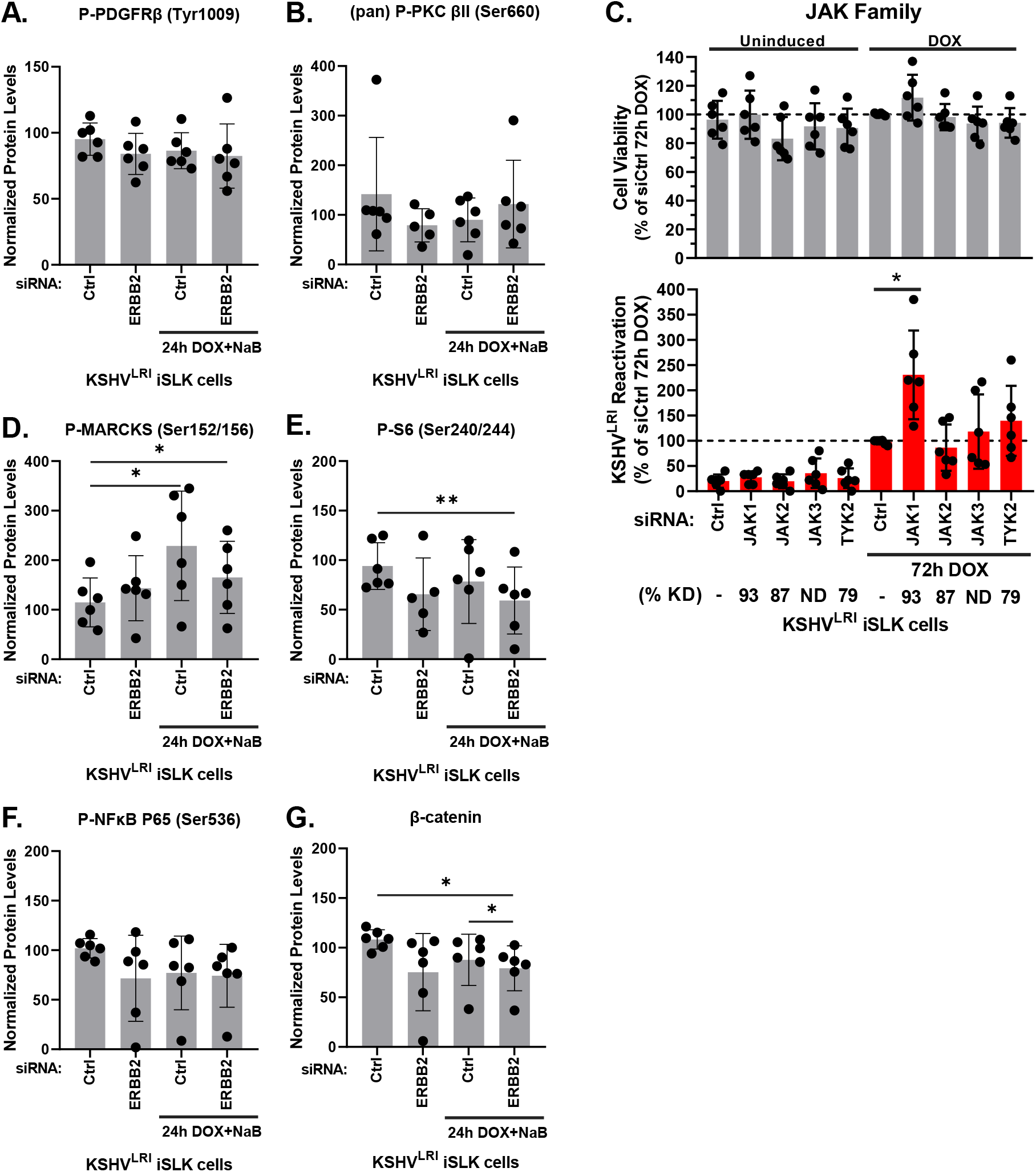
Effects of ERBB2 and reactivation on phosphorylation of downstream signaling factors. KSHV^LRI^ latently infected iSLK cells were transfected with siRNA control or siRNAs targeting ERBB2 and then 3-days later untreated or treated with DOX plus NaB for 24h. Cells were harvested, and protein lysates were analyzed using a RPPA for phosphorylation of **(A)** plasma membrane receptor PDGFRβ at Tyr^1009^ and **(B)** signaling intermediates (pan) PKC at Ser^660^. **(C)** Cell viability (grey bars) and KSHV reactivation (red bars) were measured for KSHV^LRI^ latently infected iSLK cells transfected with siRNAs targeting individual JAK family kinases and 3-days later uninduced or treated with DOX alone for 72h. Control siRNA transfected cells treated with DOX (dotted black lines) were set to 100 and data for each condition was calculated as a percent of this control. Kinase knockdown efficiencies at 3-days following siRNA transfection were determined before addition of lytic inducing drugs and graphed in Fig S6. For each knockdown, the efficiencies were averaged and listed below the corresponding kinase target as % KD. Identical to (A and B), quantification of phosphorylated **(D)** MARKS at Ser^152/156^, **(E)** S6 at Ser^240/244^, **(F)** NFκB P65 at Ser^536^, and **(G)** total protein for β-catenin was analyzed. P-values * ≤ 0.05 and ** ≤ 0.01.

**Fig S6:**
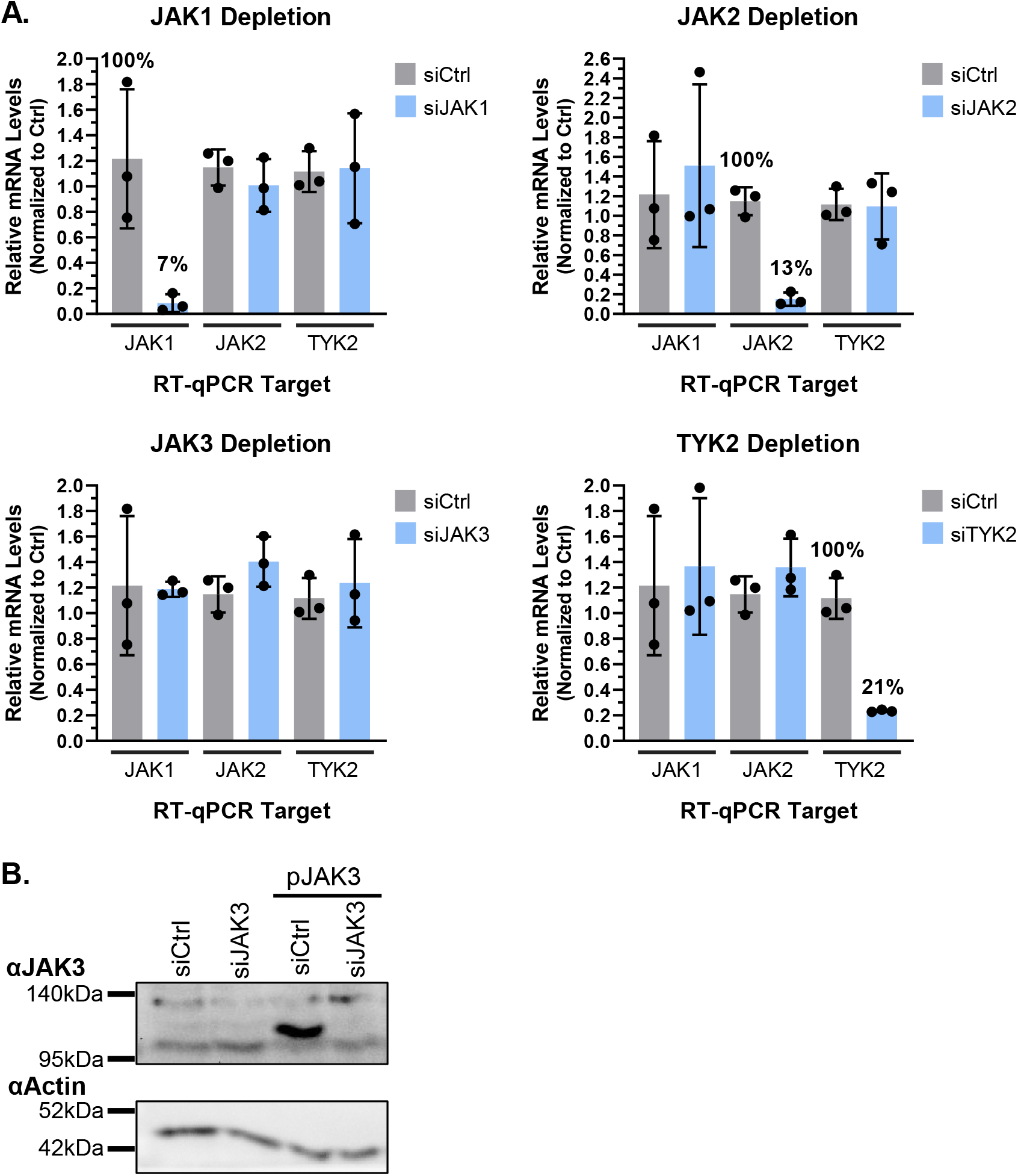
Knockdown efficiencies and specificity for JAK family members. **(A)** Knockdown specificity for siRNAs targeting JAK1, JAK2, JAK3 and TYK2 were evaluated using RT-qPCR from total RNA harvested at 3-days post transfection with siRNAs. Relative mRNA levels for JAK3 were below the level of detection for these samples. **(B)** HeLa cells were transfected with control or JAK3 targeting siRNA alone or in combination with a JAK3 expressing plasmid. Three days post transfection cells were harvested, and lysates were subjected to α-JAK3 and α-Actin immunoblotting.

### Supporting Information (SI) Captions

**S1 Table.** Kinase inhibitors list.

| **Kinase Inhibitor** | **Rank** | **Experiment** | **CAS #** | **Consortium / Vender** | **Catalog ID** |
| --- | --- | --- | --- | --- | --- |
| Baricitinib | - | Initial screen | 1187594-09-7 | NCATS | - |
| Doramapimod | - | Initial screen | 285983-48-4 | NCATS | - |
| JNK Inhibitor VIII | - | Initial screen | 894804-07-0 | NCATS | - |
| VX-11e | - | Initial screen | 896720-20-0 | NCATS | - |
| Rho Kinase Inhibitor III | - | Initial screen | 7272-84-6 | NCATS | - |
| PP121 | - | Initial screen | 1092788-83-4 | NCATS | - |
| GZD824 Dimesylate | - | Initial screen | 1421783-64-3 | NCATS | - |
| Staurosporine | - | Initial screen | 62996-74-1 | NCATS | - |
| AZD3463 | - | Initial screen | 1356962-20-3 | NCATS | - |
| NVP-BVU972 | - | Initial screen | 1185763-69-2 | NCATS | - |
| Y-33075 Dihydrochloride | - | Initial screen | 173897-44-4 | NCATS | - |
| Cobimetinib | - | Initial screen | 934660-93-2 | NCATS | - |
| WZ3146 | - | Initial screen | 1214265-56-1 | NCATS | - |
| Aminopurvalanol A | - | Initial screen | 220792-57-4 | NCATS | - |
| SB 202190 | - | Initial screen | 152121-30-7 | NCATS | - |
| PD 166285 | - | Initial screen | 212391-63-4 | NCATS | - |
| PRT062607 | - | Initial screen | 1370261-96-3 | NCATS | - |
| Akt Inhibitor VIII | - | Initial screen | 612847-09-3 | NCATS | - |
| CP-547632 | - | Initial screen | 252003-65-9 | NCATS | - |
| Bosutinib | - | Initial screen | 380843-75-4 | NCATS | - |
| AT7867 | - | Initial screen | 857531-00-1 | NCATS | - |
| Cdk1/2 Inhibitor III | - | Initial screen | 443798-55-8 | NCATS | - |
| PDK1/Akt/Flt Dual Pathway Inhibitor | - | Initial screen | 331253-86-2 | NCATS | - |
| SB 218078 | - | Initial screen | 135897-06-2 | NCATS | - |
| SCHEMBL2227980 | - | Initial screen | 1221153-14-5 | NCATS | - |
| Ruboxistaurin Mesylate | - | Initial screen | 192050-59-2 | NCATS | - |
| Cot Inhibitor-2 | - | Initial screen | 915363-56-3 | NCATS | - |
| N/A | - | Initial screen | 185039-99-0 | NCATS | - |
| Nintedanib | - | Initial screen | 928326-83-4 | NCATS | - |
| PF-477736 | 2 | Validation Set | 952021-60-2 | Sigma | PZ0186 |
| AZD3463 | 3 | Validation Set | 1356962-20-3 | Sigma | SML2590 |
| GSK-650394 | 4 | Validation Set | 890842-28-1 | MedChemExpress | HY-15192 |
| AV 412 | 5 | Validation Set | 451492-95-8 | MedChemExpress | HY-10346A |
| ASP-3026 | 6 | Validation Set | 1097917-15-1 | MedChemExpress | HY-13326 |
| Gefitinib | 7 | Validation Set | 184475-35-2 | Sigma | SML1657 |
| Lestaurtinib | 8 | Validation Set | 111358-88-4 | Sigma | C7869 |
| Afatinib | 9 | Validation Set | 850140-72-6 | MedChemExpress | HY-10261 |
| K252a | 14 | Validation Set | 99533-80-9 | Abcam | AB120419 |
| PKR Inhibitor | 18 | Validation Set | 608512-97-6 | Calbiochem/Sigma | 527451 |
| IRAK1/4 Inhibitor | 140 | Validation Set | 509093-47-4 | Sigma | I5409 |
| JAK3 Inhibitor VI | 205 | Validation Set | 856436-16-3 | Sigma | 420126 |
| Tofacitinib | - | JAK Family | 540737-29-9 | LC Laboratories | T-1377 |

**S2 Table.**
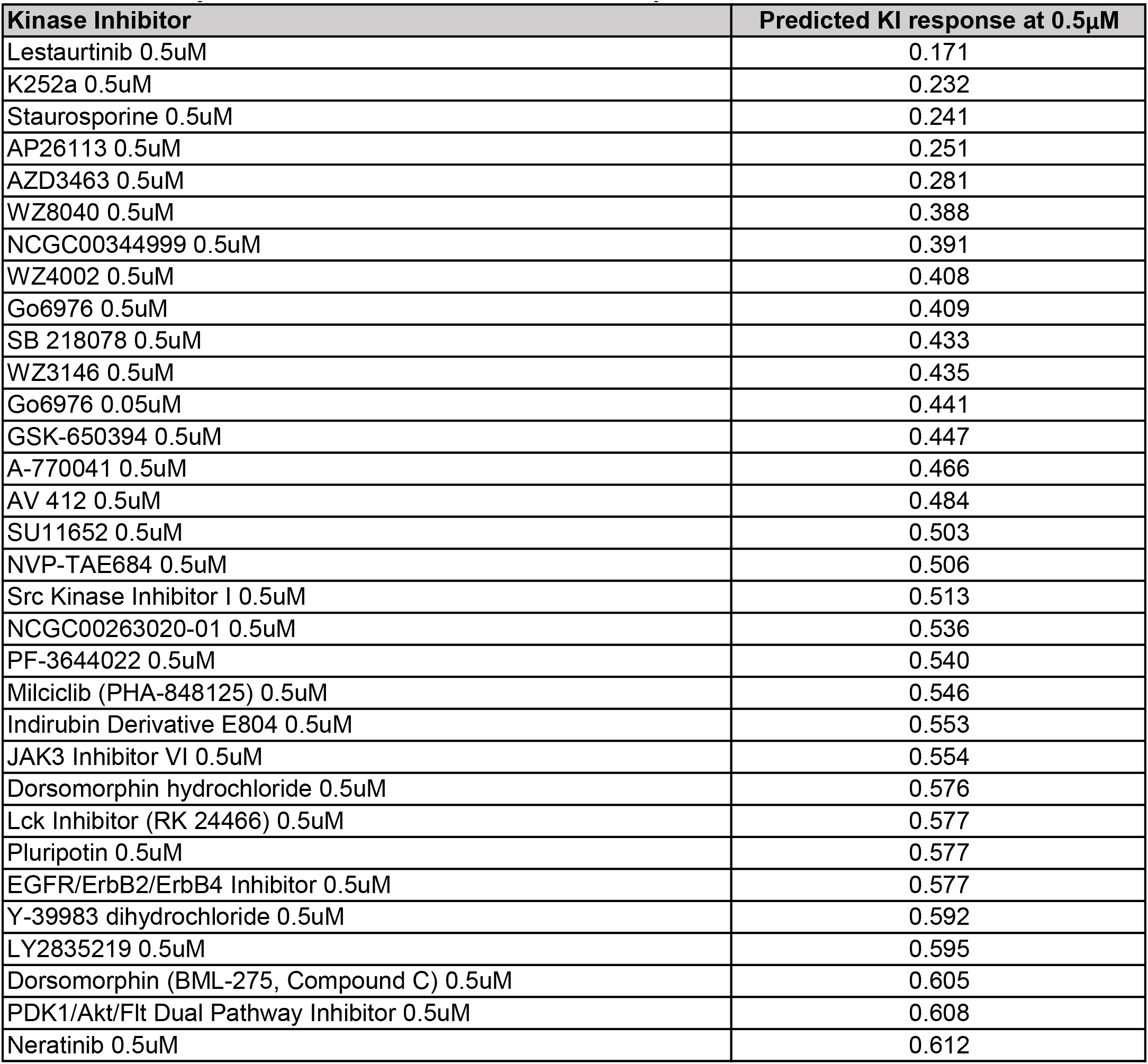
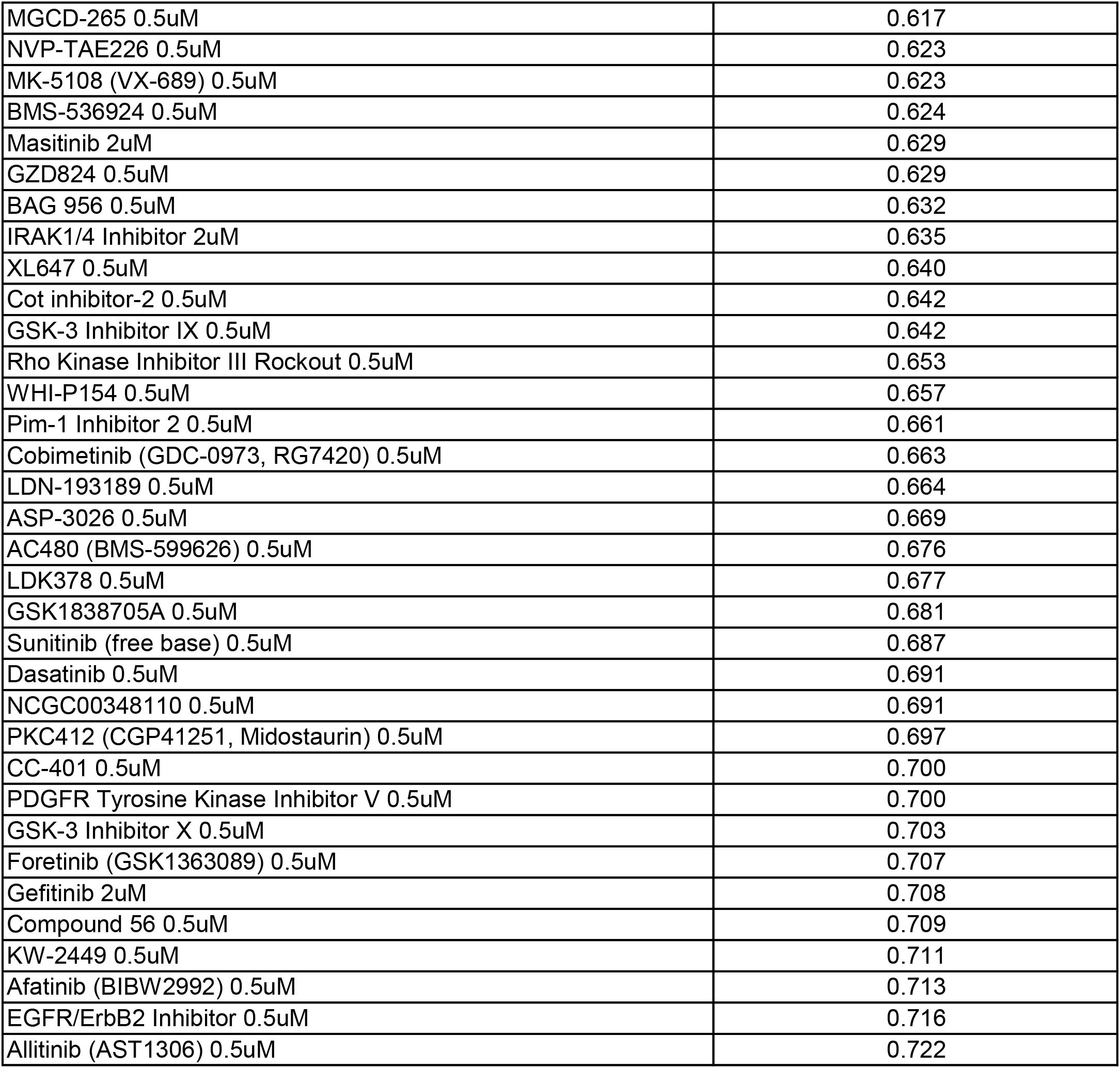
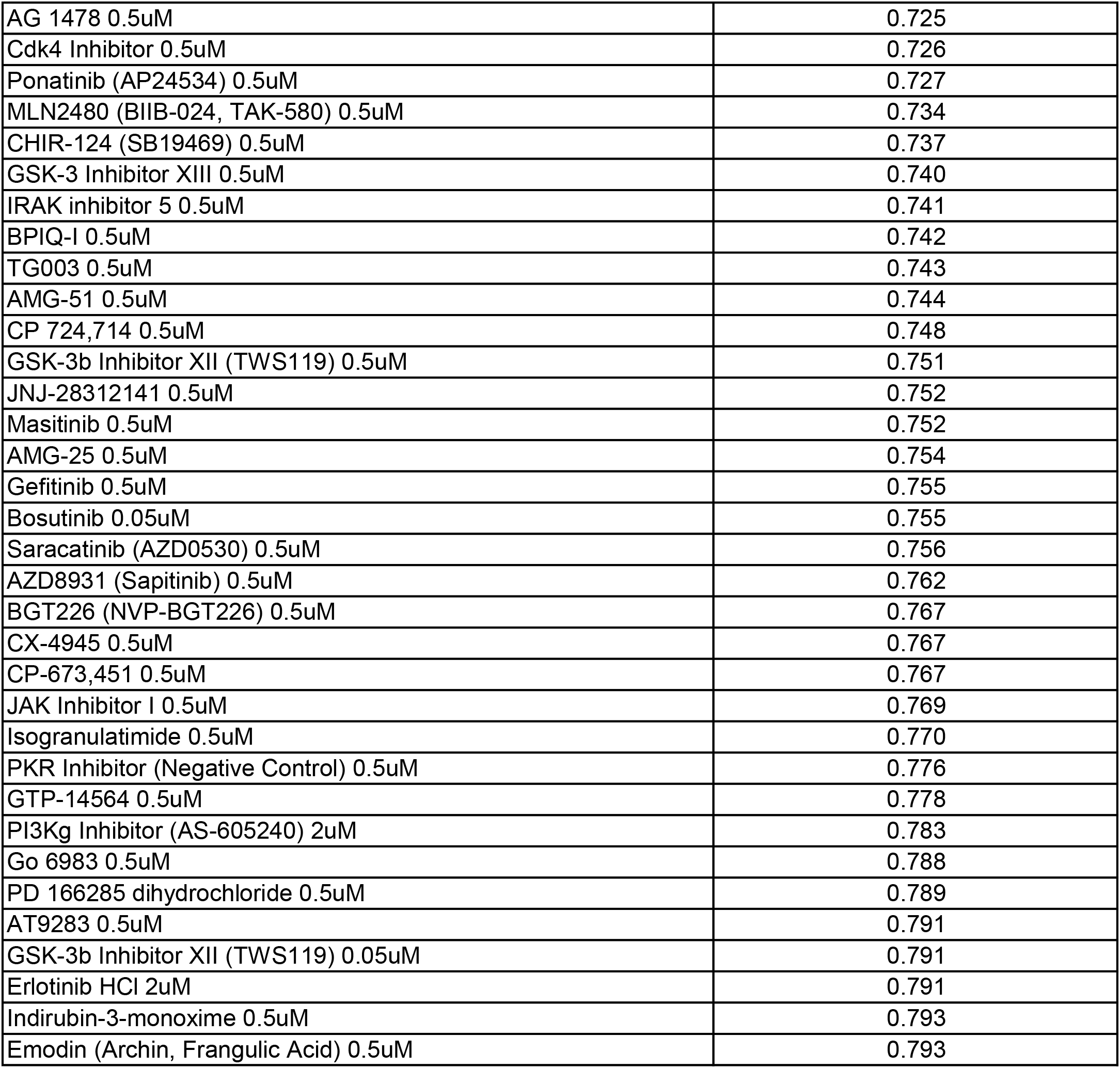
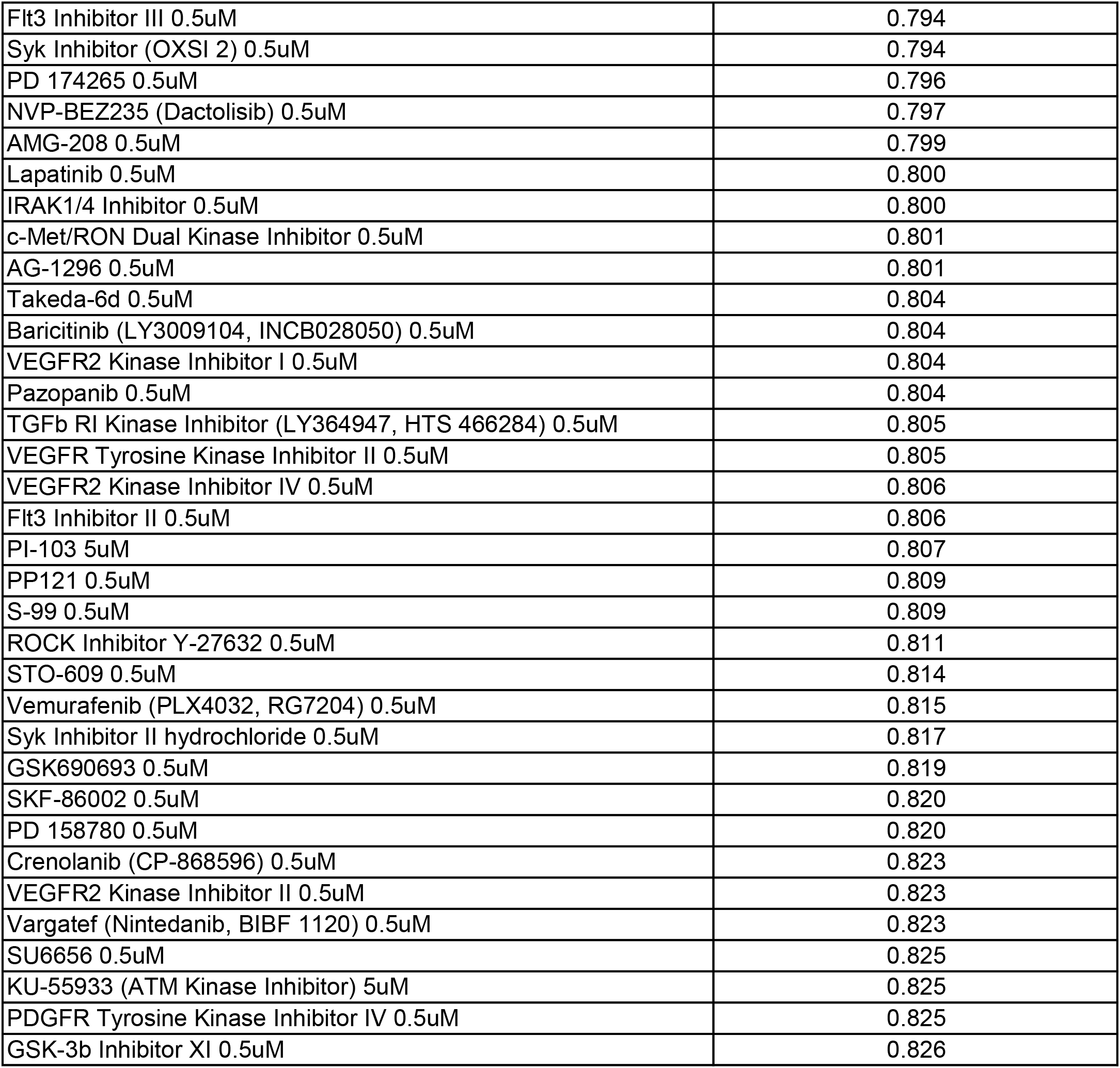
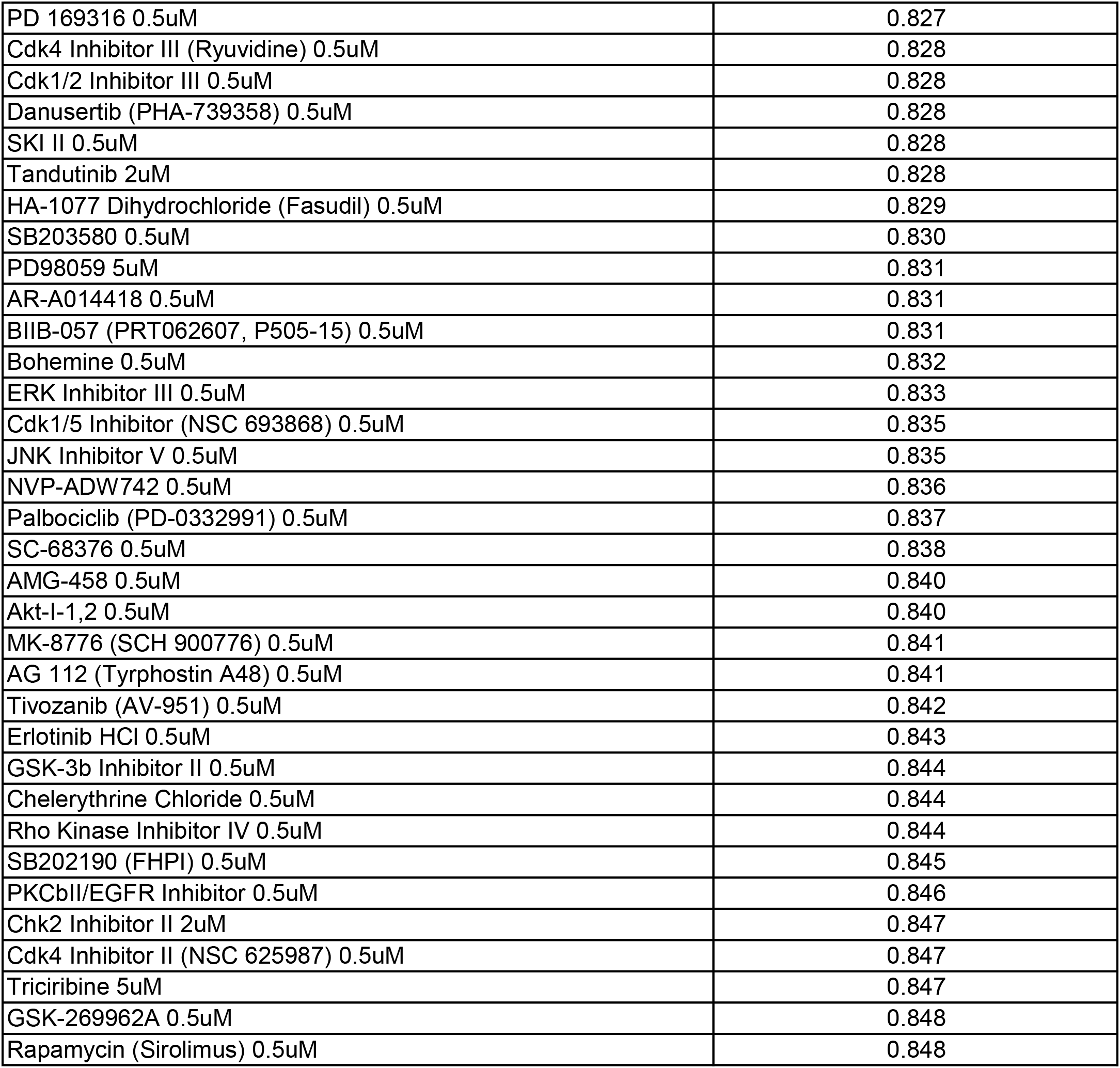
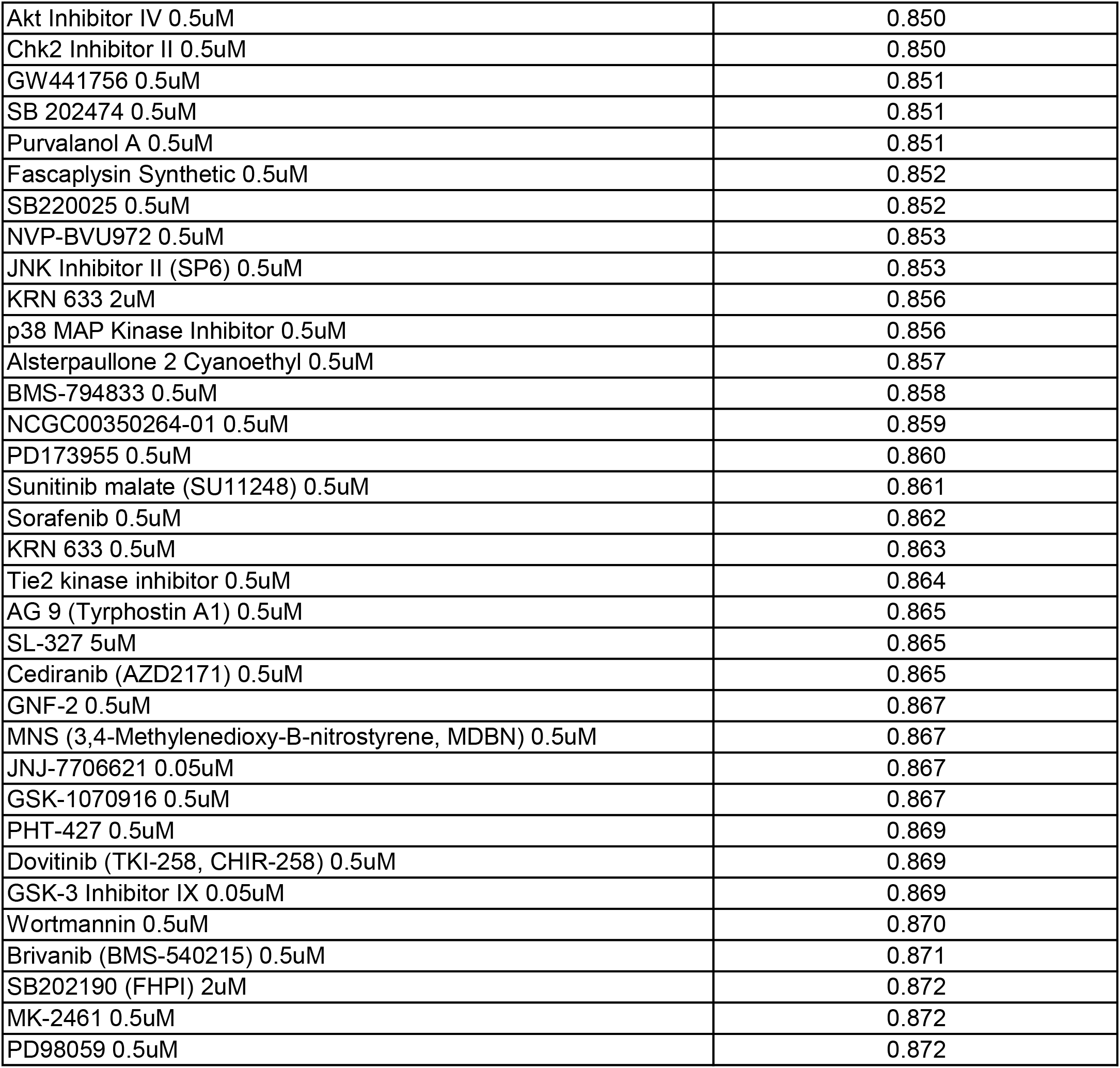
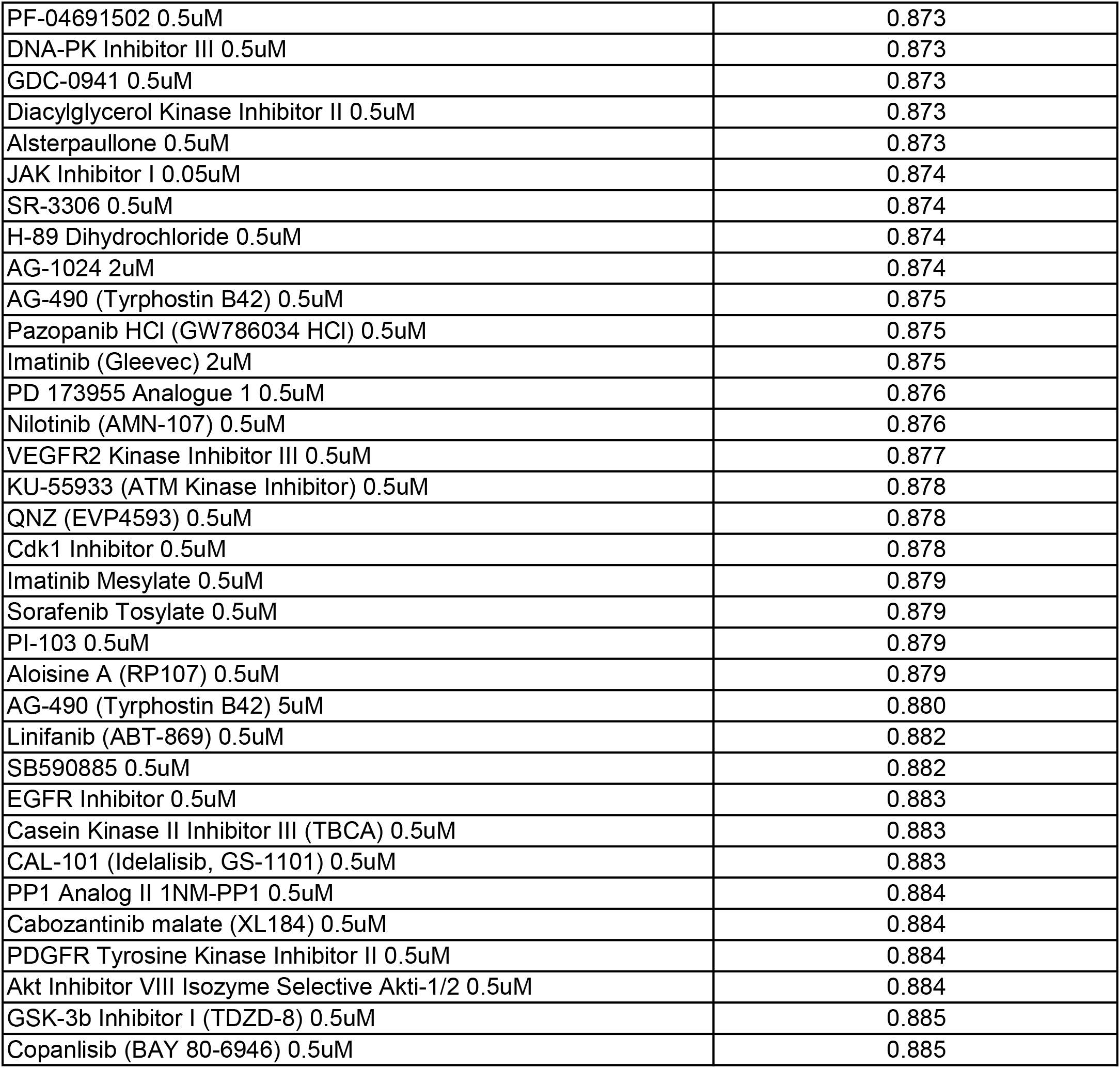
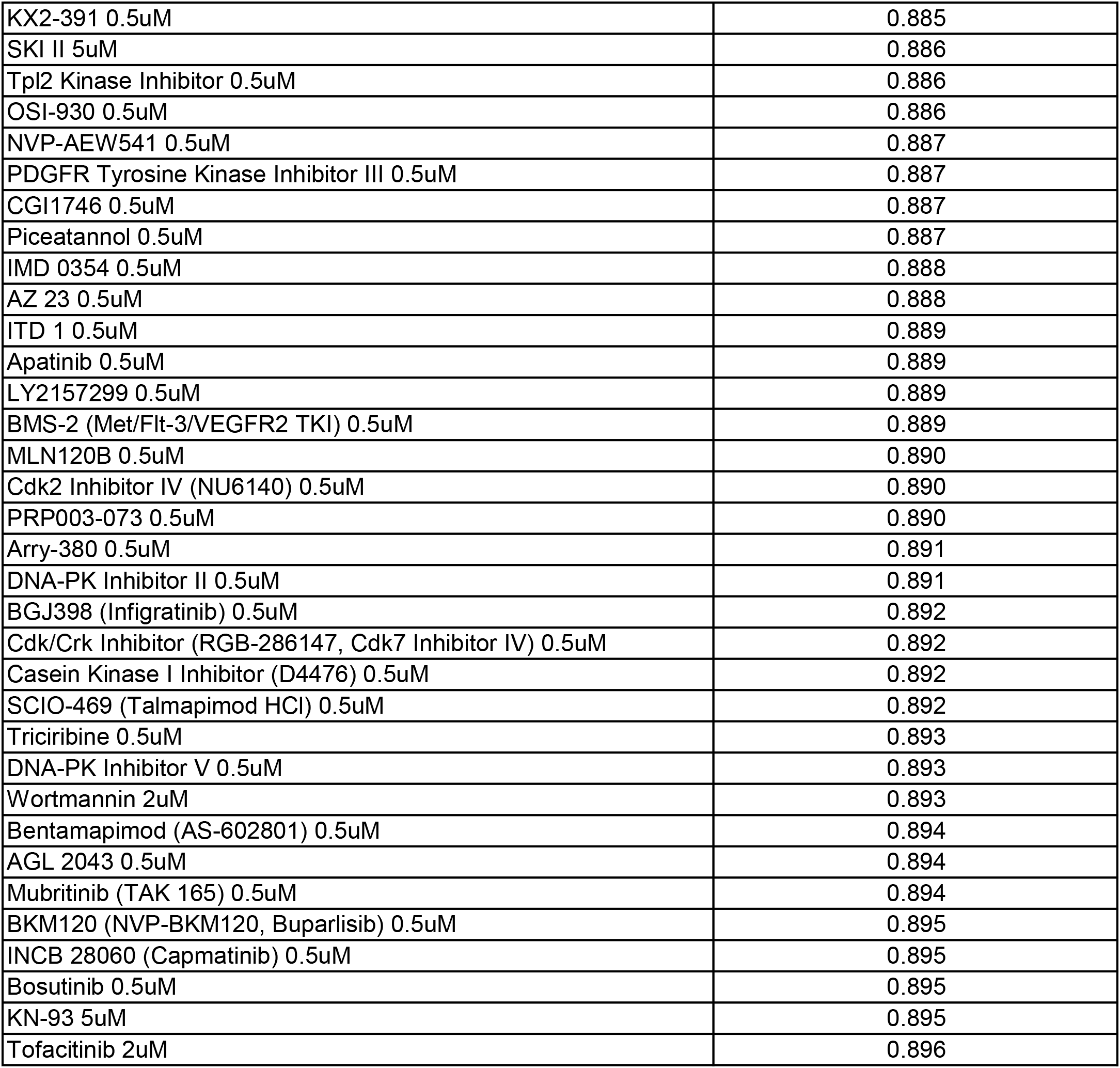
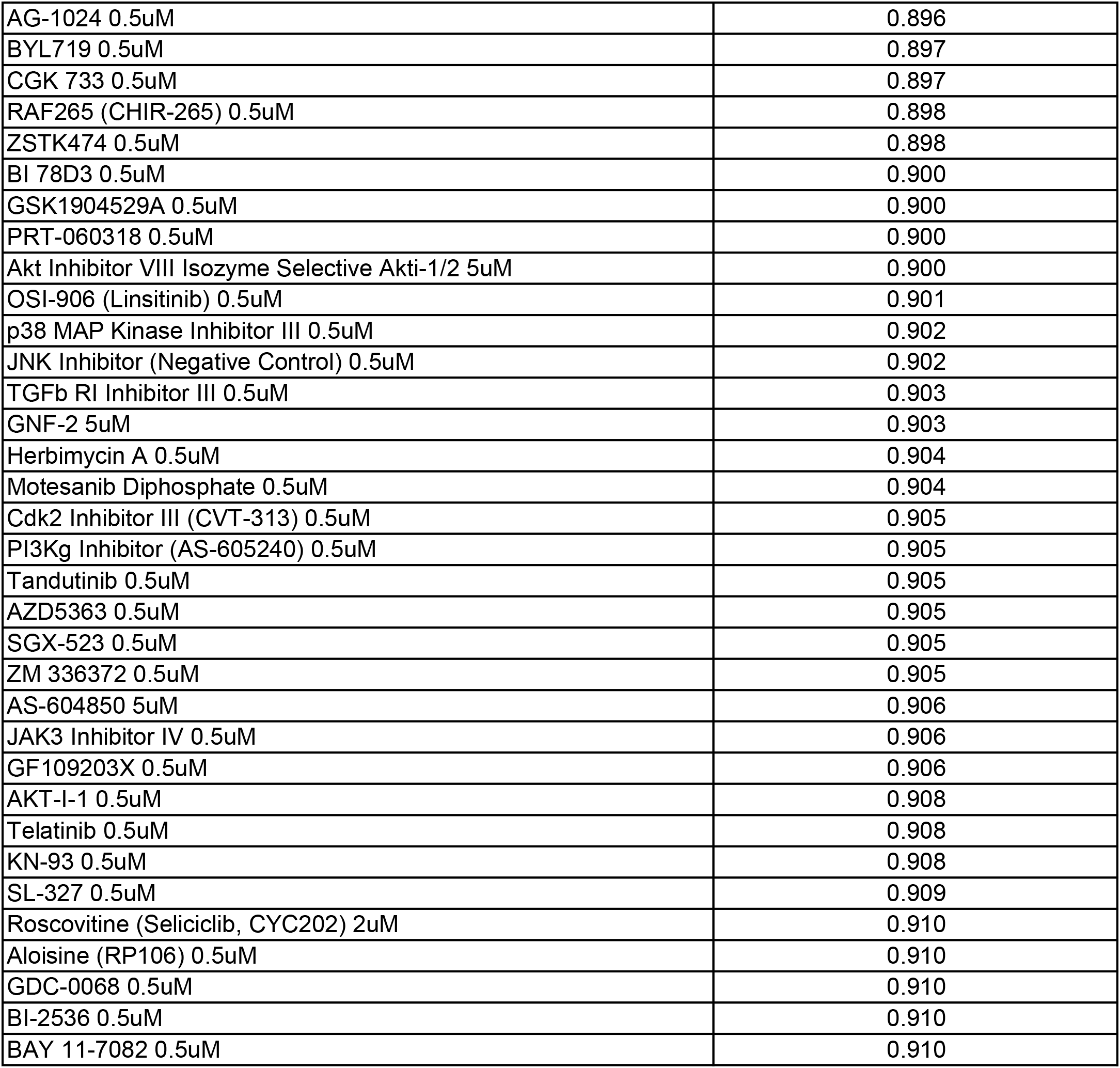
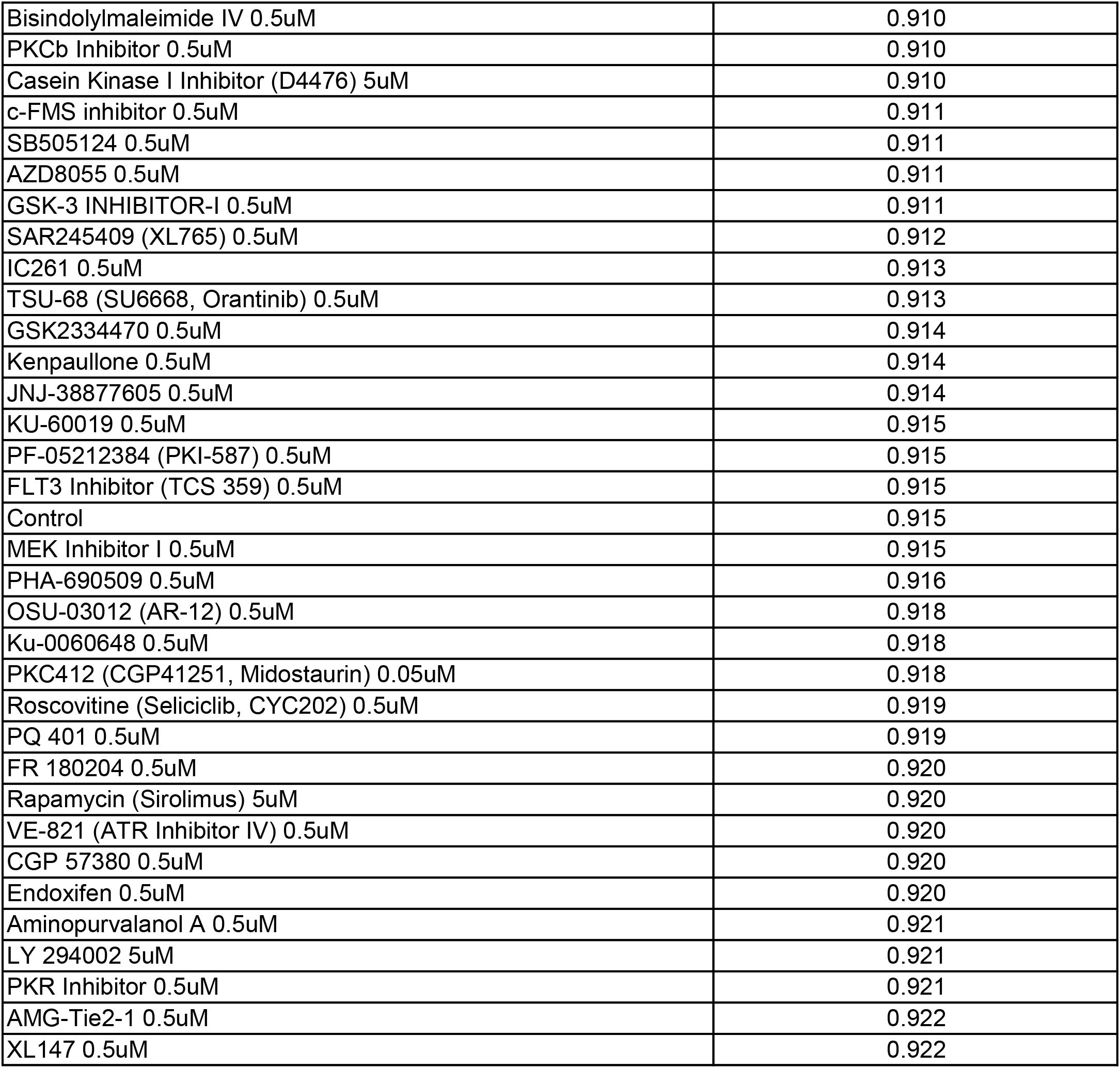
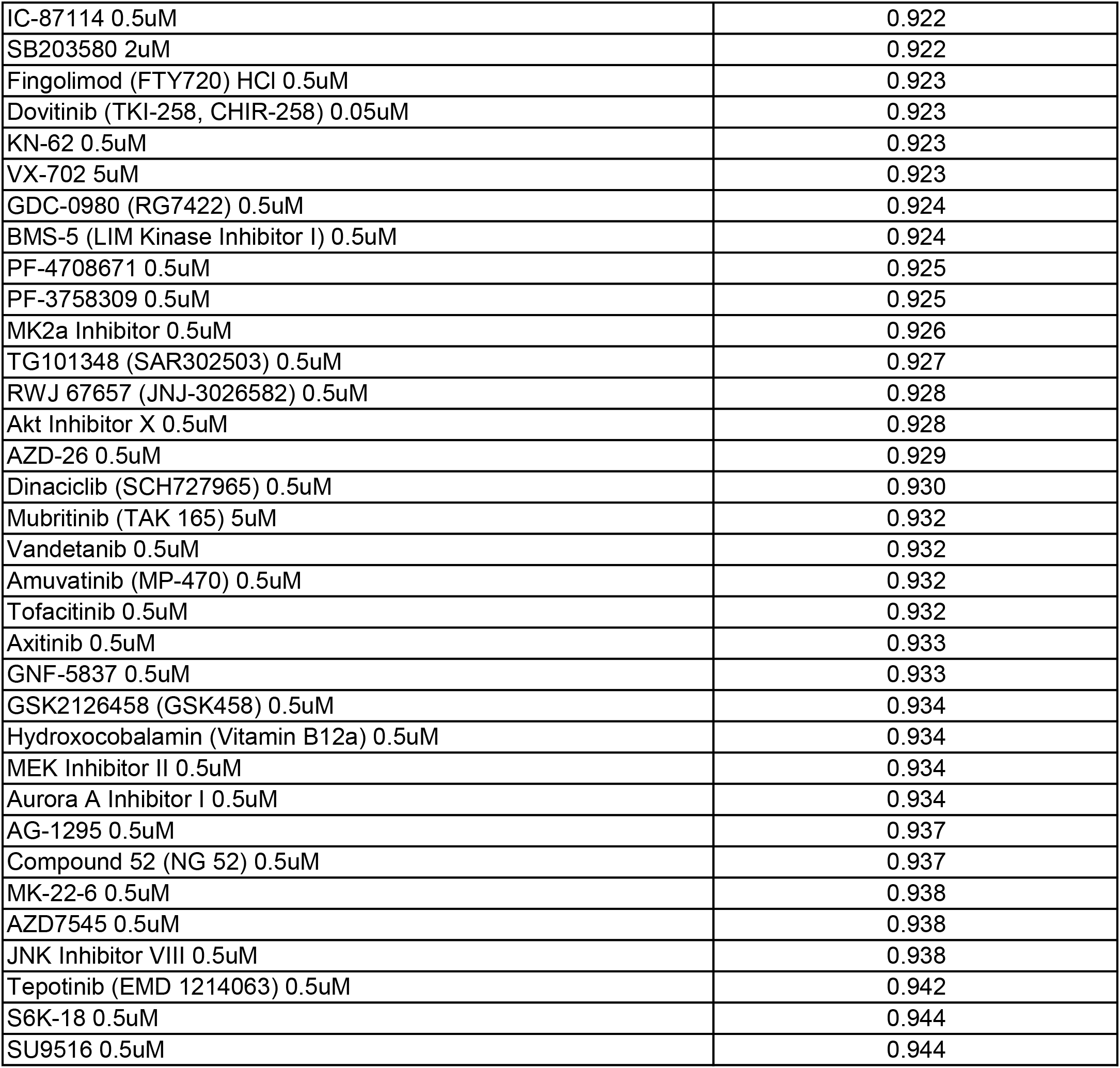
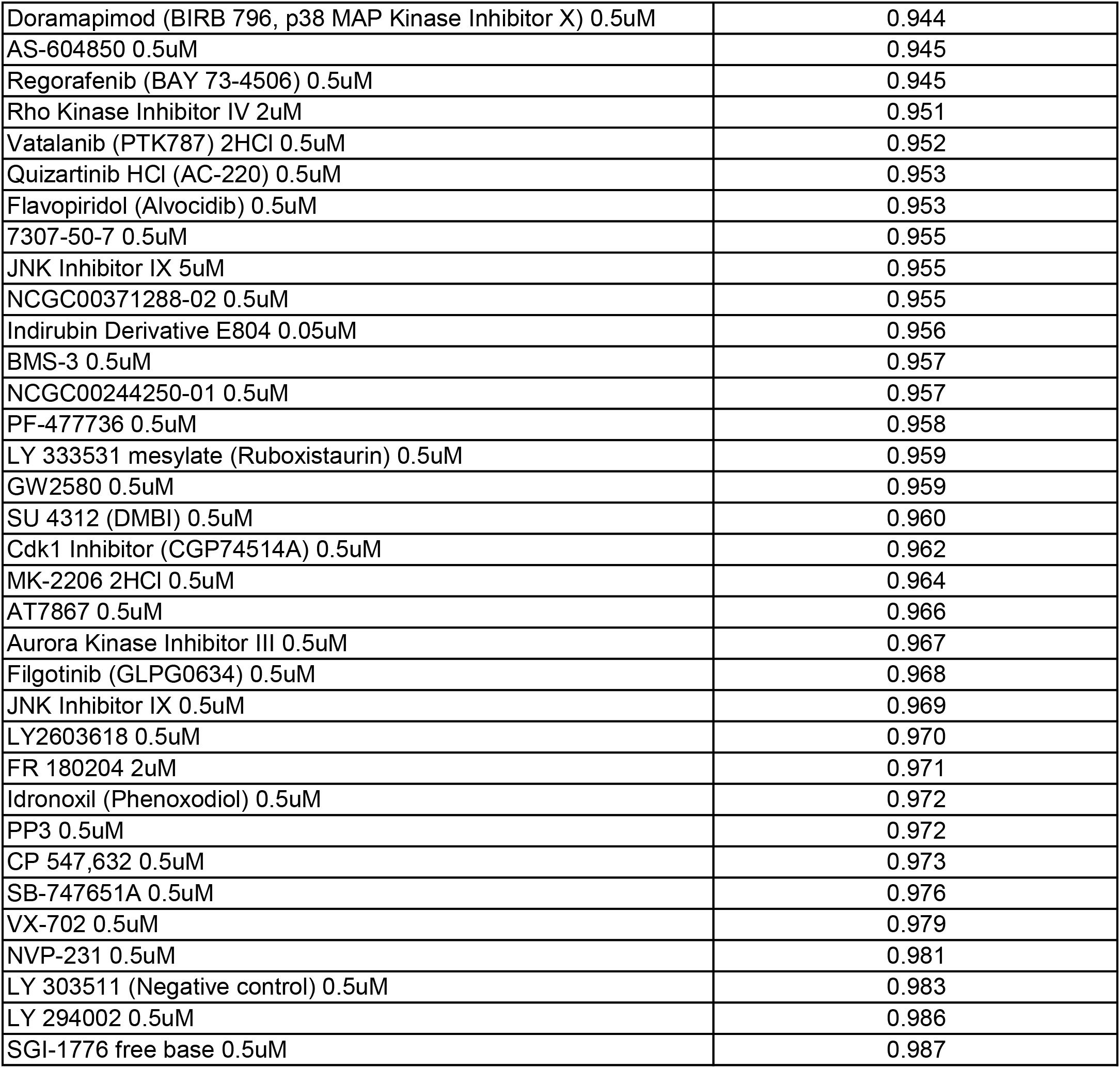
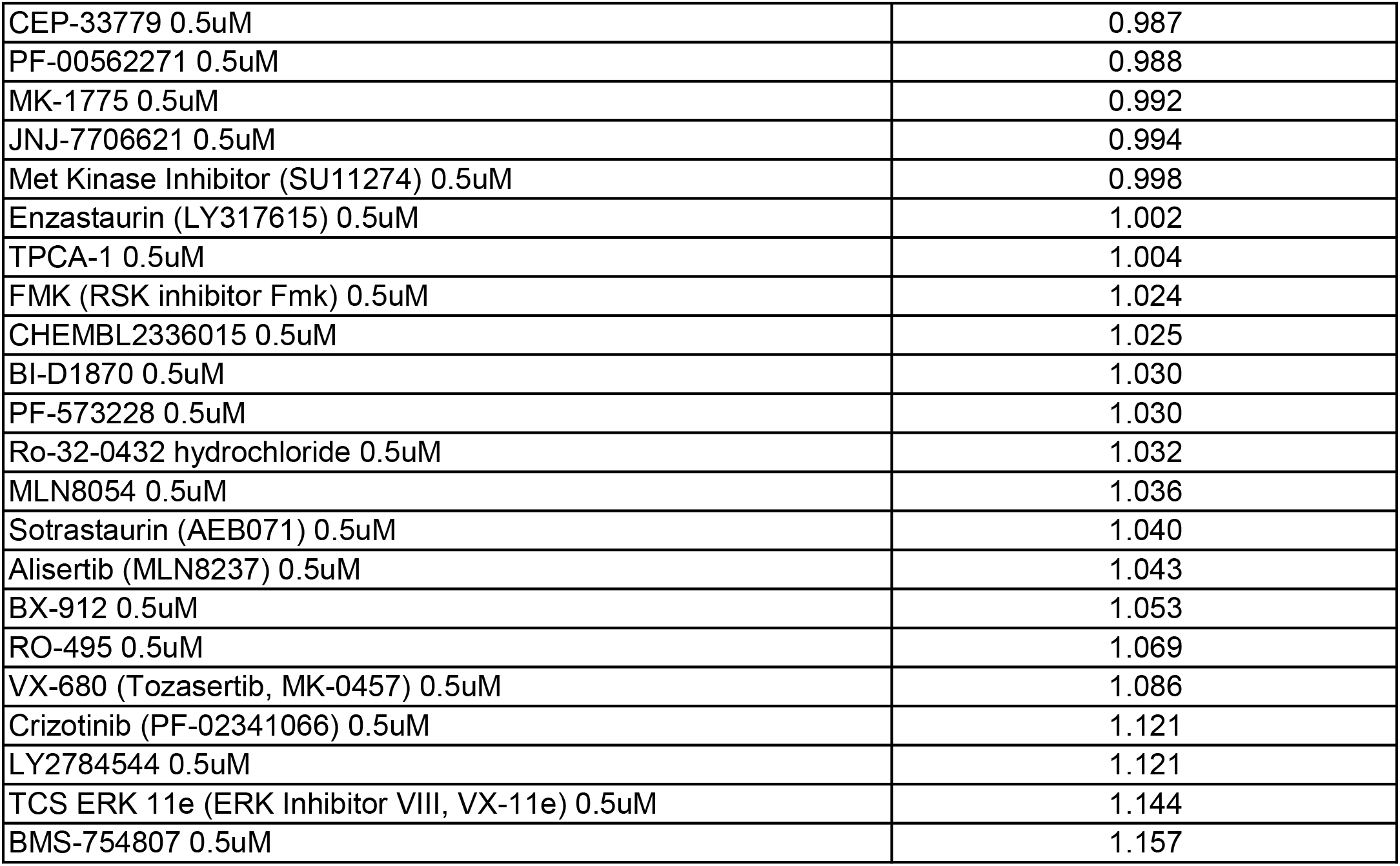
DOX plus NaB condition KiR screen final predicted kinase inhibitors.

**S3 Table.** DOX plus NaB condition KiR screen final predicted kinases.

| Number | Kinase | Average Coefficient |
| --- | --- | --- |
| 1 | ITK | 0.00247 |
| 2 | *PKG2 (PRKG2)\** | 0.00223 |
| 3 | MAP3K8 (COT1) | 0.00210 |
| 4 | FRK | 0.00183 |
| 5 | CLK1 | 0.00191 |
| 6 | ERBB4 (HER4) | 0.00168 |
| 7 | MAP2K2 (MEK2) | 0.00125 |
| 8 | CAMK2G | 0.00083 |
| 9 | MKNK2 (MNK2) | 0.00086 |
| 10 | PBK | 0.00079 |
| 11 | TEC | 0.00072 |
| 12 | LRRK2 | 0.00055 |
| 13 | MAP4K4 (HGK) | 0.00045 |
| 14 | DSTYK (RIPK5) | 0.00066 |

*\* No expression data for iSLK cells containing latent or lytic replicating virus [34]. Excluded from validated kinases.*

**S4 Table.**
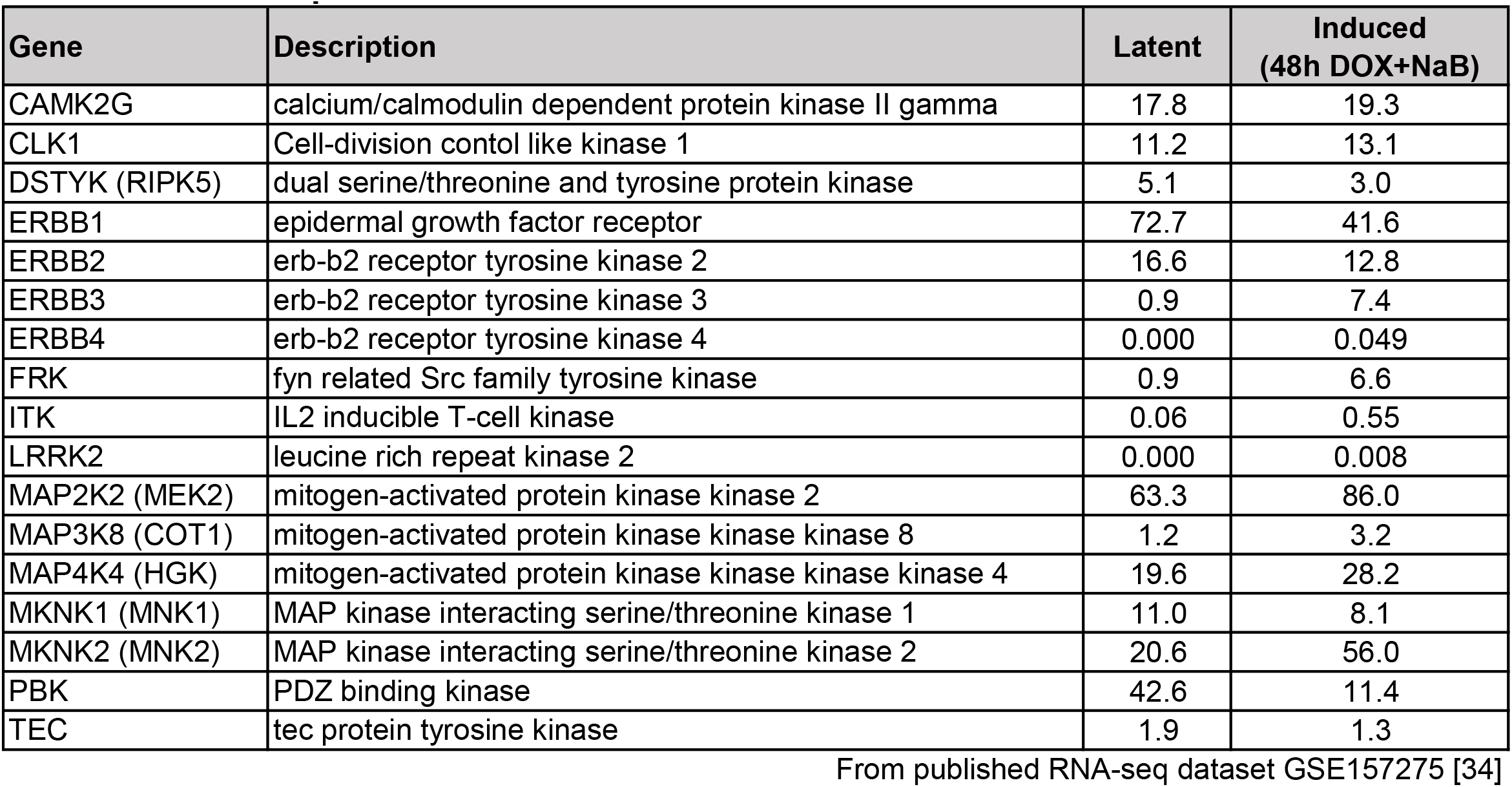
Kinase expression data from KSHV BAC16 infected iSLK cells.

The gene expression data normalized as fragments per kilobase of exon per million mapped fragments (FPKM) were taken from the published RNA-seq dataset GSE157275 [34] for the kinases predicted from the kinome screen under lytic induction and the additional ERBB and MKNK family members. These data were organized into a table showing the normalized kinase expression data from BAC16 latently infected iSLK cells and lytically induced cells at 48h post treatment with 50 µg/ml DOX plus 1.2 mM NaB.

**S5 Table.**
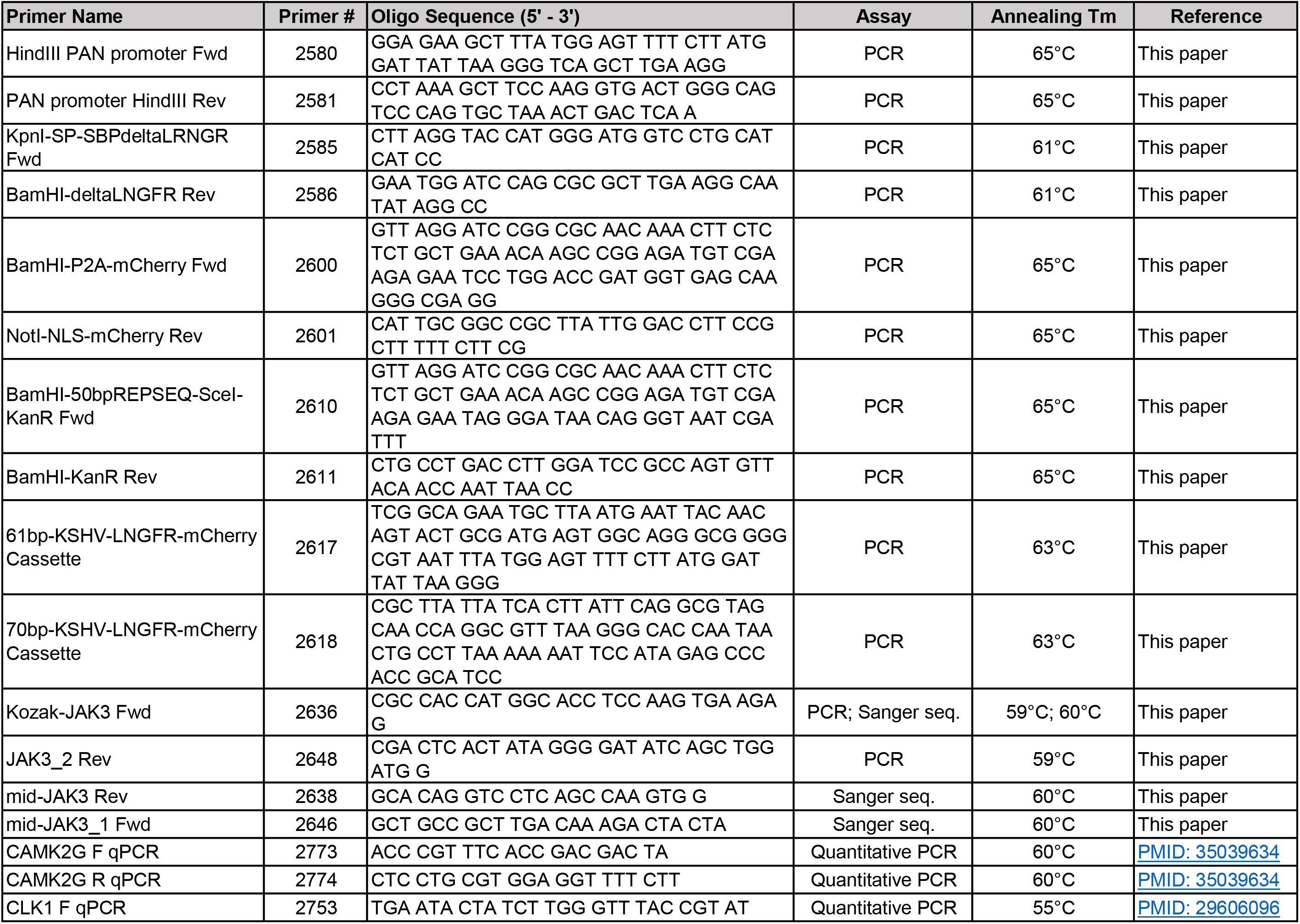
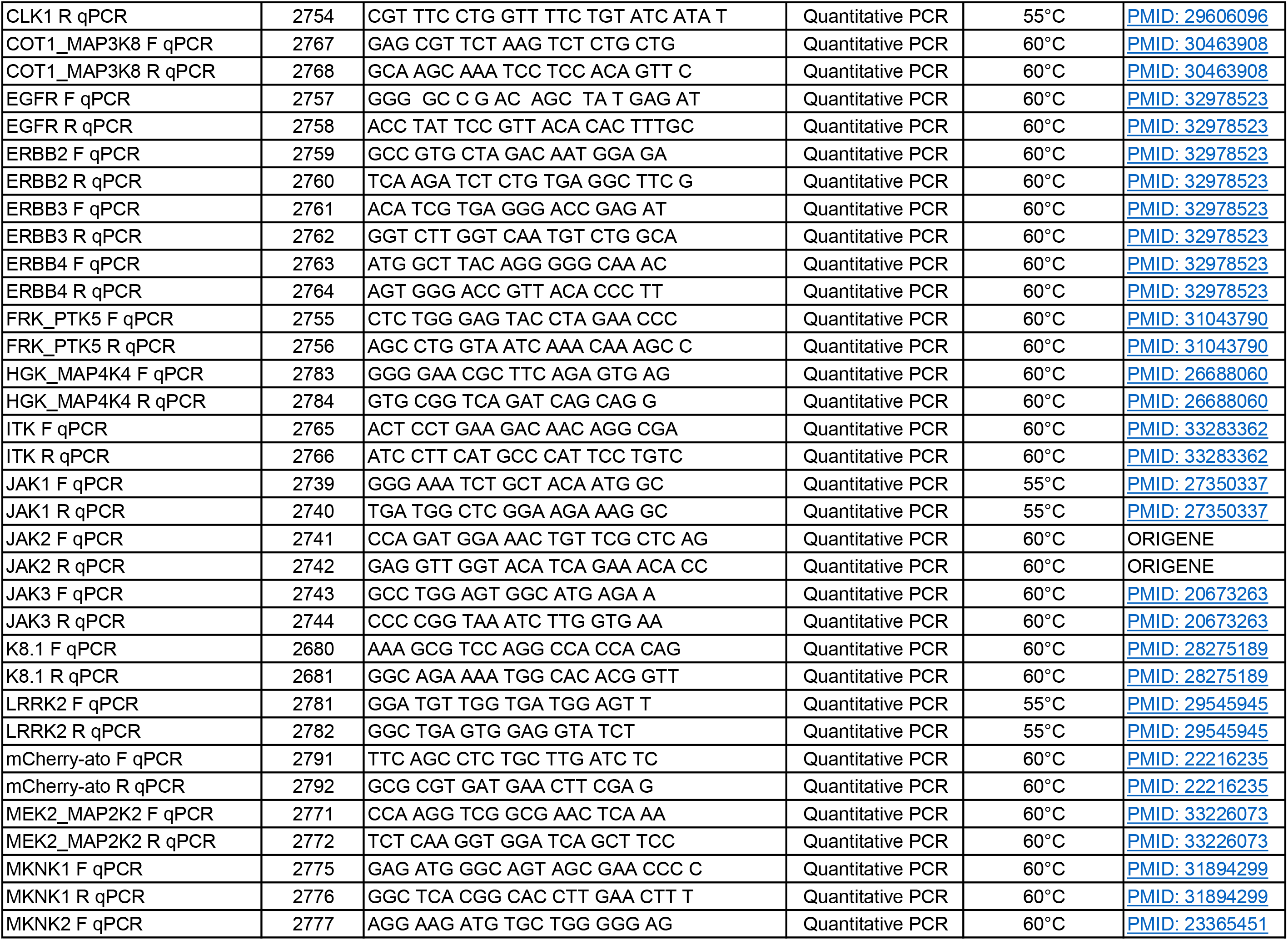
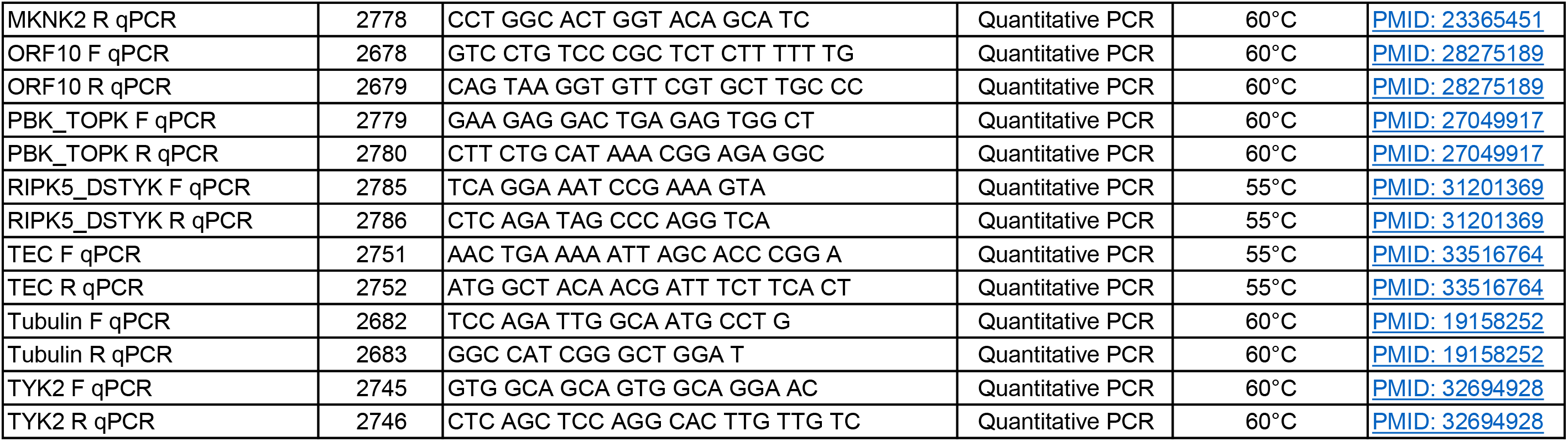
Primers.

**S6 Table.** Dharmacon siRNA target and ID.

| siRNA Target | Dharmacon Catalog ID |
| --- | --- |
| CAMK2G | L-004536-00-0005 |
| CLK1 | L-004800-00-0005 |
| DSTYK | L-004049-00-0005 |
| ERBB1 | L-003114-00-0005 |
| ERBB2 | L-003126-00-0005 |
| ERBB3 | L-003127-00-0005 |
| ERBB4 | L-003128-00-0005 |
| FRK | L-003139-00-0005 |
| ITK | L-003144-00-0005 |
| JAK1 | L-003145-00-0005 |
| JAK2 | L-003146-00-0005 |
| JAK3 | L-003147-00-0005 |
| LRRK2 | L-006323-00-0005 |
| MAP2K2 | L-003573-00-0005 |
| MAP3K8 | L-003511-00-0005 |
| MAP4K4 | L-003971-00-0005 |
| MKNK1 | L-004879-00-0005 |
| MKNK2 | L-004908-00-0005 |
| Non-targeting control | D-001810-10-05 |
| PBK | L-005390-00-0005 |
| TEC | L-003177-00-0005 |
| TYK2 | L-003182-00-0005 |

**S7 Table.** Antibodies

| Antibody Name | Phospho-site | Vender / Source | Catalog ID / Reference | Assay |
| --- | --- | --- | --- | --- |
| Actin | N/A | Sigma | A2066 | Immunoblot |
| β-actin | N/A | Sigma | A1978 | Protein microarray |
| Her2 c-terminus (ERBB2) | N/A | Rabbit derived | Child et al. 1999;<br>PMID: 10216954 | Immunoblot |
| JAK3 (B-12) | N/A | Santa Cruz | sc-6932 | Immunoblot |
| LANA | N/A | Rabbit derived | Lagunoff et al. 2002;<br>PMID: 11836422;<br>a kind gift from A. Polson<br>and D. Ganem | Immunoblot;<br>Immunofluorescence |
| p75 NGF receptor (LNGFR) | N/A | Abcam | ab52987 | Immunoblot |
| P-AKT | Ser473 | Cell Signaling | 4058 | Protein microarray |
| P-CREB1 | Ser133 | Cell Signaling | 9198 | Protein microarray |
| P-EGFR (ERBB1) | Tyr1173 | Cell Signaling | 4407 | Protein microarray |
| P-MARCKS | er152/156 | Cell Signaling | 2741 | Protein microarray |
| P-MET | Tyr1349 | Cell Signaling | 3133 | Protein microarray |
| P-NFκB P65 | Ser536 | Cell Signaling | 3033 | Protein microarray |
| P-P44/42 MAPK (ERK1/2) | Thr202/Tyr204 | Cell Signaling | 4377 | Protein microarray |
| P-PDGFRβ | Tyr1009 | Cell Signaling | 3124 | Protein microarray |
| P-PKC (pan) βII | Ser660 | Cell Signaling | 9371 | Protein microarray |
| P-S6 ribosomal protein | Ser240/244 | Cell Signaling | 2215 | Protein microarray |
| P-SRC | Tyr416 | Cell Signaling | 2101 | Protein microarray |
| P-STAT1 | Tyr701 | Cell Signaling | 9167 | Protein microarray |
| P-STAT3 | Tyr705 | Cell Signaling | 9145 | Protein microarray |
| Streptavidin, Alexa Fluor 680 conjugate | N/A | Invitrogen | S21378 | Immunofluorescence |
| β-catenin | N/A | Cell Signaling | 9582 | Protein microarray |
| Goat αRabbit, Alexa Fluor 488 | N/A | Invitrogen | A11008 | Immunofluorescence |
| Goat αRabbit, Alkaline Phosphatase | N/A | Invitrogen | T2191 | Immunoblot |
| Goat αMouse, IRDye 800CW | N/A | LI-COR | 926-32210 | Protein microarray |
| Goat αRabbit, IRDye 680LT | N/A | LI-COR | 925-68071 | Protein microarray |

